# Clathrin adaptors mediate two sequential pathways of intra-Golgi recycling

**DOI:** 10.1101/2021.03.17.435835

**Authors:** Jason C. Casler, Adam H. Krahn, Areti Pantazopoulou, Natalie Johnson, Kasey J. Day, Benjamin S. Glick

**Affiliations:** Department of Molecular Genetics and Cell Biology, The University of Chicago, 920 East 58th St., Chicago, IL 60615, USA

## Abstract

The pathways of membrane traffic within the Golgi apparatus are not fully known. This question was addressed using the yeast *Saccharomyces cerevisiae*, in which the maturation of individual Golgi cisternae can be visualized. We recently proposed that the AP-1 clathrin adaptor mediates intra-Golgi recycling late in the process of cisternal maturation. Here, we demonstrate that AP-1 cooperates with the Ent5 clathrin adaptor to recycle a set of Golgi transmembrane proteins. This recycling can be detected by removing AP-1 and Ent5, thereby diverting the AP-1/Ent5-dependent Golgi proteins into an alternative recycling loop that involves traffic to the plasma membrane followed by endocytosis. Unexpectedly, various AP-1/Ent5-dependent Golgi proteins show either intermediate or late kinetics of residence in maturing cisternae. We infer that the AP-1/Ent5 pair mediates two sequential intra-Golgi recycling pathways that define two classes of Golgi proteins. This insight can explain the polarized distribution of transmembrane proteins in the Golgi.

## Introduction

The structure and composition of the Golgi apparatus are well described. In many cell types, including animal and plant cells, the Golgi consists of stacks of disk-like cisternae (Farquhar and Palade, 1981). In other organisms such as the budding yeast *Saccharomyces cerevisiae*, the cisternae are not stacked (Mowbrey and Dacks, 2009; Papanikou and Glick, 2009). Despite these differences in organization, the basic functions of the Golgi are conserved. Newly synthesized cargo proteins arrive at the Golgi from the endoplasmic reticulum (ER), advance from early to late cisternae, and ultimately depart from the *trans*-Golgi network (TGN) to either the plasma membrane or the endosomal/lysosomal/vacuolar system (De Matteis and Luini, 2008). Golgi resident transmembrane proteins include components involved in membrane traffic as well as enzymes involved in the glycosylation and proteolytic processing of cargo molecules (Banfield, 2011). On the surface of Golgi cisternae are peripheral membrane proteins that include GTPases, vesicle coat proteins, and vesicle tethers. These peripheral membrane proteins cooperate with lipids and transmembrane proteins to drive traffic to, from, and within the Golgi (Munro, 2002; Gillingham and Munro, 2016). The molecular components of the Golgi are extensively characterized in biochemical and structural terms.

Less is known about the broader operating principles of the Golgi machine. Increasing evidence favors a cisternal maturation model in which Golgi cisternae assemble *de novo* from ER-derived membranes, progressively mature, and ultimately disintegrate at the TGN stage by forming secretory vesicles and other carriers (Glick and Luini, 2011). Golgi maturation can be visualized directly in *S. cerevisiae* by labeling resident Golgi proteins with fluorescent tags and then observing the arrival and departure of the tagged proteins in individual cisternae (Glick and Nakano, 2009). Although the dynamic properties of the Golgi are still debated for mammalian cells (Patterson et al., 2008; Pfeffer, 2010; Pellett et al., 2013), a variety of experimental findings combined with the strong conservation of membrane traffic components support the generality of the maturation process (Glick and Nakano, 2009).

A challenge for the maturation model is to explain Golgi polarity, which is a central feature of this organelle. In a stacked Golgi, some resident transmembrane proteins are concentrated at the *cis* side of the stack whereas other resident transmembrane proteins are concentrated in the middle of the stack or at the *trans* side (Dunphy and Rothman, 1985; Rabouille et al., 1995; Tie et al., 2016). A similar polarized distribution is seen in the non-stacked Golgi of *S. cerevisiae* (Kim et al., 2016; Day et al., 2018; Tojima et al., 2019). These observations led to the view that the Golgi consists of compartments designated *cis*, medial/*trans*, and TGN (Dunphy and Rothman, 1985; Mellman and Simons, 1992). According to the maturation model, each *cis* cisterna matures into a medial/*trans* cisterna and then into a TGN cisterna, while resident Golgi proteins constantly recycle from older to younger cisternae. We originally proposed that Golgi compartmentation could be reconciled with the maturation model if a given compartment represents a kinetic stage of maturation (Day et al., 2013). Such a kinetic stage would begin when a recycling pathway delivers a subset of resident Golgi proteins to a maturing cisterna, and would end when a recycling pathway removes those proteins. However, that concept proved unwieldy because Golgi compartments have never been rigorously defined and because Golgi recycling pathways show temporal overlap.

As an alternative, we suggest that the Golgi should be seen not as a set of compartments, but rather as a set of maturing cisternae controlled by a molecular logic circuit that switches various membrane traffic pathways on and off in a particular sequence (Pantazopoulou and Glick, 2019). To characterize this logic circuit, an important step is to clarify which membrane recycling pathways operate at the Golgi, and when. COPI-coated vesicles have been shown to mediate Golgi-to-ER recycling and intra-Golgi recycling of certain transmembrane proteins (Rabouille and Klumperman, 2005; Barlowe and Miller, 2013). In yeast, COPI is present during approximately the first half of the maturation process and seems to act selectively in the recycling of early Golgi proteins (Papanikou et al., 2015; Ishii et al., 2016; Kim et al., 2016). We have proposed that a later-acting Golgi recycling pathway involves clathrin-coated vesicles that form with the aid of the AP-1 adaptor (Papanikou et al., 2015; Day et al., 2018; Casler and Glick, 2019). Yeast AP-1 has been implicated in the recycling of transmembrane TGN proteins (Valdivia et al., 2002; Foote and Nothwehr, 2006; Liu et al., 2008; Spang, 2015), and we found that AP-1 is restricted to the TGN, implying that AP-1 mediates an intra-Golgi recycling pathway downstream of COPI (Day et al., 2018). Mammalian AP-1 also has a role in retrograde traffic (Hinners and Tooze, 2003; Hirst et al., 2012), so this recycling function of AP-1 may be conserved.

Yeast AP-1 interacts with the epsin-related clathrin adaptor Ent5, which has homology to mammalian epsinR (Duncan et al., 2003; Costaguta et al., 2006; Copic et al., 2007). The relationship between AP-1 and Ent5 is complex and still poorly understood, but the data suggest that these two adaptors have partially overlapping functions and can act independently (Costaguta et al., 2006; Copic et al., 2007). It therefore seems likely that AP-1 and Ent5 cooperate to recycle Golgi transmembrane proteins at the TGN.

Although recycling of Golgi transmembrane proteins is probably crucial for cell growth, yeast strains lacking both AP-1 and Ent5 grow robustly (Costaguta et al., 2006). A possible explanation is that in the absence of AP-1 and Ent5, some Golgi transmembrane proteins escape to the plasma membrane and then recycle by endocytosis (Valdivia et al., 2002; Liu et al., 2008). Such a bypass mechanism for AP-1/Ent5-dependent proteins is easy to picture because the yeast Golgi also serves as an early endosome, with endocytic vesicles fusing around the time that an early Golgi cisterna matures into a TGN cisterna (Day et al., 2018). We now demonstrate that simultaneous removal of AP-1 and Ent5 does indeed cause a set of Golgi transmembrane proteins to recycle via the plasma membrane. As a control, proteins that either reside in the early Golgi or recycle from prevacuolar endosome (PVE) compartments are unaffected by removal of AP-1/Ent5. These results confirm the existence of AP-1/Ent5-dependent intra-Golgi recycling.

If multiple Golgi transmembrane proteins follow the same recycling pathway, they should show similar kinetics of arrival and departure during cisternal maturation (Pantazopoulou and Glick, 2019). We were therefore surprised to find that AP-1/Ent5-dependent Golgi proteins fall into two kinetic classes. A class of proteins that reside in the Golgi during a late phase of maturation have previously been designated TGN residents, and a class of proteins that reside in the Golgi during an intermediate phase of maturation have been designated medial/*trans* residents. Apparently, the AP-1/Ent5 pair mediates two sequential pathways of intra-Golgi recycling. This finding provides a mechanistic explanation for how various Golgi transmembrane proteins are concentrated in different cisternae.

## Results

### The AP-1/Ent5 pair recycles membrane during TGN maturation

Payne and colleagues reported that AP-1 and Ent5 showed similar kinetic signatures late in the process of yeast TGN maturation, although the data hinted at some differences between the two adaptors (Daboussi et al., 2012). To obtain a clear view of AP-1 and Ent5 kinetics, we devised a procedure for smoothing and averaging the noisy fluorescence traces obtained from analyzing individual cisternae by 4D confocal microscopy (see Methods). Figure 1A shows frames from Video S1, in which a yeast strain expressed the TGN marker Sec7-mScarlet together with the AP-1 subunit Apl2-GFP plus Ent5-HaloTag coupled to the far-red dye JFX_646_ (Losev et al., 2006; Casler et al., 2019; Grimm et al., 2021). A representative cisterna was chosen for analysis. In accord with prior results (Day et al., 2018; Casler et al., 2019), AP-1 arrived about halfway through the Sec7 time course and departed after Sec7. Figure 1B shows quantification of the fluorescence signals from this cisterna. In Figure 1C, the traces from 18 such events were smoothed and averaged. The results indicate that for a typical cisterna, Ent5 arrives a few seconds before AP-1, accumulates to its maximal level faster than AP-1, and begins to depart while AP-1 levels are still rising. Thus, AP-1 and Ent5 overlap substantially during TGN maturation but show distinct kinetic signatures. The AP-1/Ent5 pair is a candidate for mediating membrane recycling from maturing TGN cisternae.

**Figure 1.**
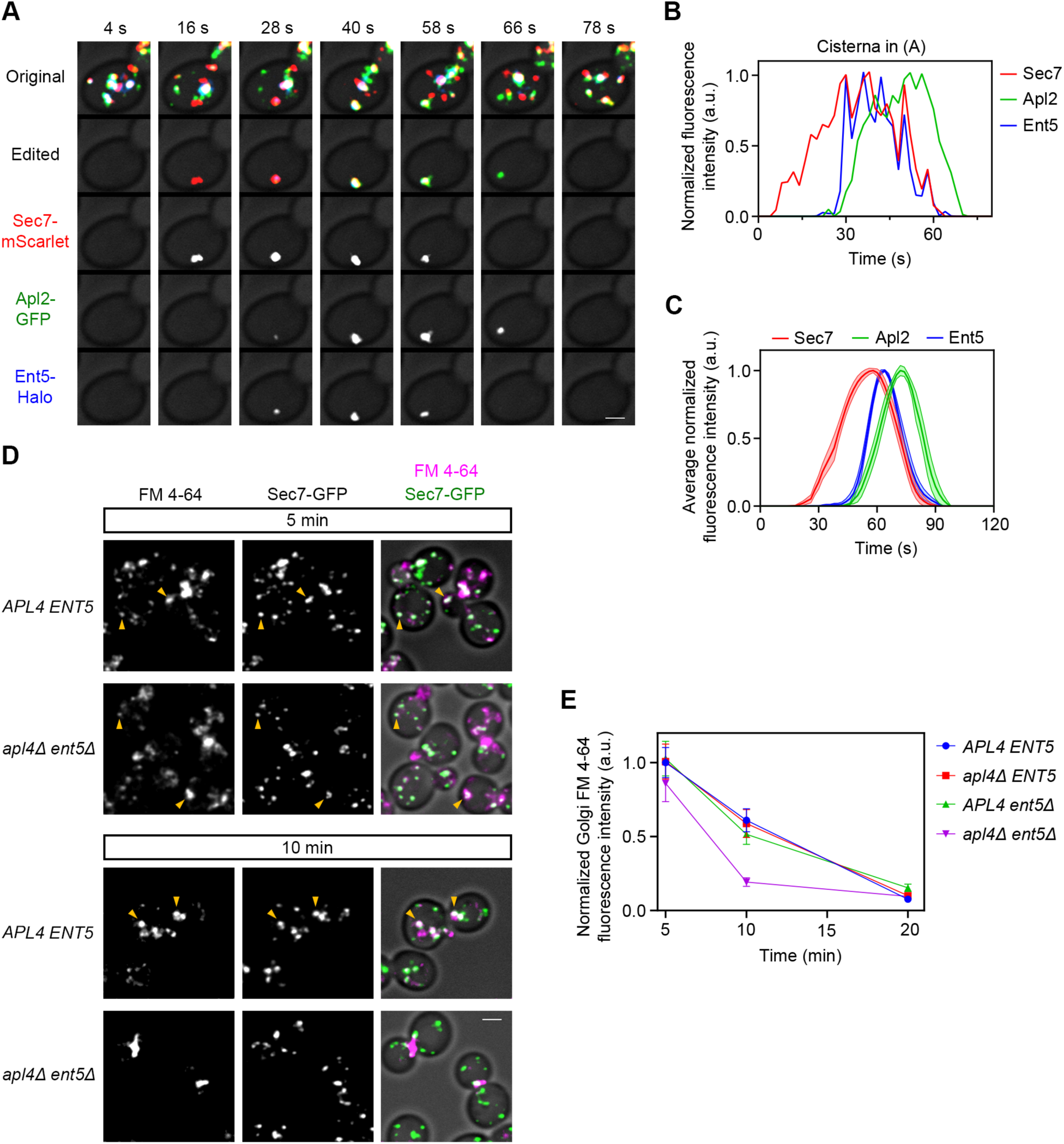
AP-1 and Ent5 Operate at the TGN and Are Responsible for the Persistent TGN Localization of Internalized FM 4-64. See also Video S1. (A) Maturation kinetics of AP-1 and Ent5 compared to Sec7. A strain expressing the TGN marker Sec7-mScarlet, the AP-1 subunit Apl2-GFP, and the clathrin adaptor Ent5-HaloTag was grown to mid-log phase, labeled with JFX dye, and imaged by 4D confocal microscopy. Shown are average projected Z-stacks at the indicated time points from Video S1. The upper row shows the complete projections, the second row shows edited projections that include only the cisterna being tracked, and the subsequent rows show the individual fluorescence channels from the edited projections. Scale bar, 2 µm. (B) Quantification of tagged Golgi proteins during a typical maturation event. Depicted are the normalized fluorescence intensities in arbitrary units (a.u.) for the cisterna tracked in (A). (C) Smoothed and averaged traces showing the relative kinetic signatures of Sec7, Apl2, and Ent5. Data were obtained for 18 representative cisternae. Lines show mean values, and shaded areas show 95% confidence intervals. (D) Comparison of internalized FM 4-64 in *APL4 ENT5* and *apl4Δ ent5Δ* strains. Cells expressing Sec7-GFP were grown to mid-log phase and incubated with FM 4-64FX (a fixable version of FM 4-64) during a 3-min pulse, followed by a chase with the quencher SCAS. Representative images are shown from the 5- and 10-min time points during the chase. Individual fluorescence channels are shown in grayscale with merged images on the right. Arrowheads mark examples of TGN structures that contain FM 4-64. Scale bar, 2 µm. (E) Quantification of the analysis in (D). For the indicated time points, the Sec7-GFP signals were used to create masks to measure the TGN-associated FM 4-64 fluorescence in *APL4 ENT*5, *apl4Δ ENT5*, *APL4 ent5Δ*, and *apl4Δ ent5Δ* strains. Plotted are the average TGN-associated FM 4-64 signals, normalized to the value in wild-type cells at 5 min. Error bars represent SEM. At least 100 cells of each strain were analyzed per time point.

To test this hypothesis, we revisited an apparent paradox: when the bulk membrane marker dye FM 4-64 was internalized by endocytosis to the yeast TGN, dye signal persisted in the TGN for many minutes even though TGN cisternae turn over on a much faster time scale (Day et al., 2018). Our proposed explanation was that membrane components continually recycle from older to younger TGN cisternae (Day et al., 2018). We therefore predicted that removal of AP-1/Ent5 would reduce the persistence of FM 4-64 in the TGN. Figure 1D shows internalized FM 4-64 after a 3-min pulse followed by a 5- or 10-min chase, in cells expressing Sec7-GFP. In a wild-type strain, FM 4-64 was visible in many of the Sec7-containing cisternae at both time points. By contrast, in a strain lacking both Ent5 and the AP-1 subunit Apl4, FM 4-64 was readily detected in Sec7-containing cisternae only at the 5-min time point. At the 10-min time point in the *apl4Δ ent5Δ* strain, FM 4-64 was often seen at sites of polarized secretion (Finger and Novick, 1998) such as the bud necks of large-budded cells (Figure 1D), suggesting that recycling of FM 4-64 to the cell surface (Wiederkehr et al., 2000) was accelerated in this mutant. Figure 1E quantifies FM 4-64 colocalization with Sec7 in the wild-type and double mutant strains and also in the two single mutants. As predicted, FM 4-64 persistence in the TGN was dramatically reduced in the double mutant. These results suggest that AP-1/Ent5 recycles membrane during TGN maturation.

### A bypass pathway of recycling from the plasma membrane permits Golgi operation in the absence of AP-1/Ent5

If the AP-1/Ent5 pair plays a major role in intra-Golgi recycling, how do cells survive in the absence of these adaptors? A likely explanation is that in an *apl4Δ ent5Δ* strain, transmembrane proteins that would normally recycle within the Golgi travel instead to the plasma membrane and then recycle to the Golgi by endocytosis (Figure 2A). This concept fits with reports that certain TGN proteins could be accumulated at the plasma membrane in AP-1 mutants by blocking endocytosis with the actin polymerization inhibitor latrunculin A (Valdivia et al., 2002; Liu et al., 2008). As a more specific alternative to latrunculin A, we chose CK-666, which inhibits the Arp2/3 complex and prevents endocytosis at actin patches (Hetrick et al., 2013; Burke et al., 2014; Antkowiak et al., 2019). To ensure that CK-666 could act at full potency, the transcription factors Pdr1 and Pdr3 were deleted to prevent expression of pleiotropic drug transporters (Schüller et al., 2007; Barrero et al., 2016). A control experiment confirmed that within minutes after addition, CK-666 redistributed the actin patch component Abp1 (Goode et al., 2001; Huckaba et al., 2004) to the cytosol and completely blocked endocytic internalization of FM 4-64 (Figure 2B,C). We therefore predicted that treatment of an *apl4Δ ent5Δ* strain with CK-666 would perturb Golgi function.

**Figure 2.**
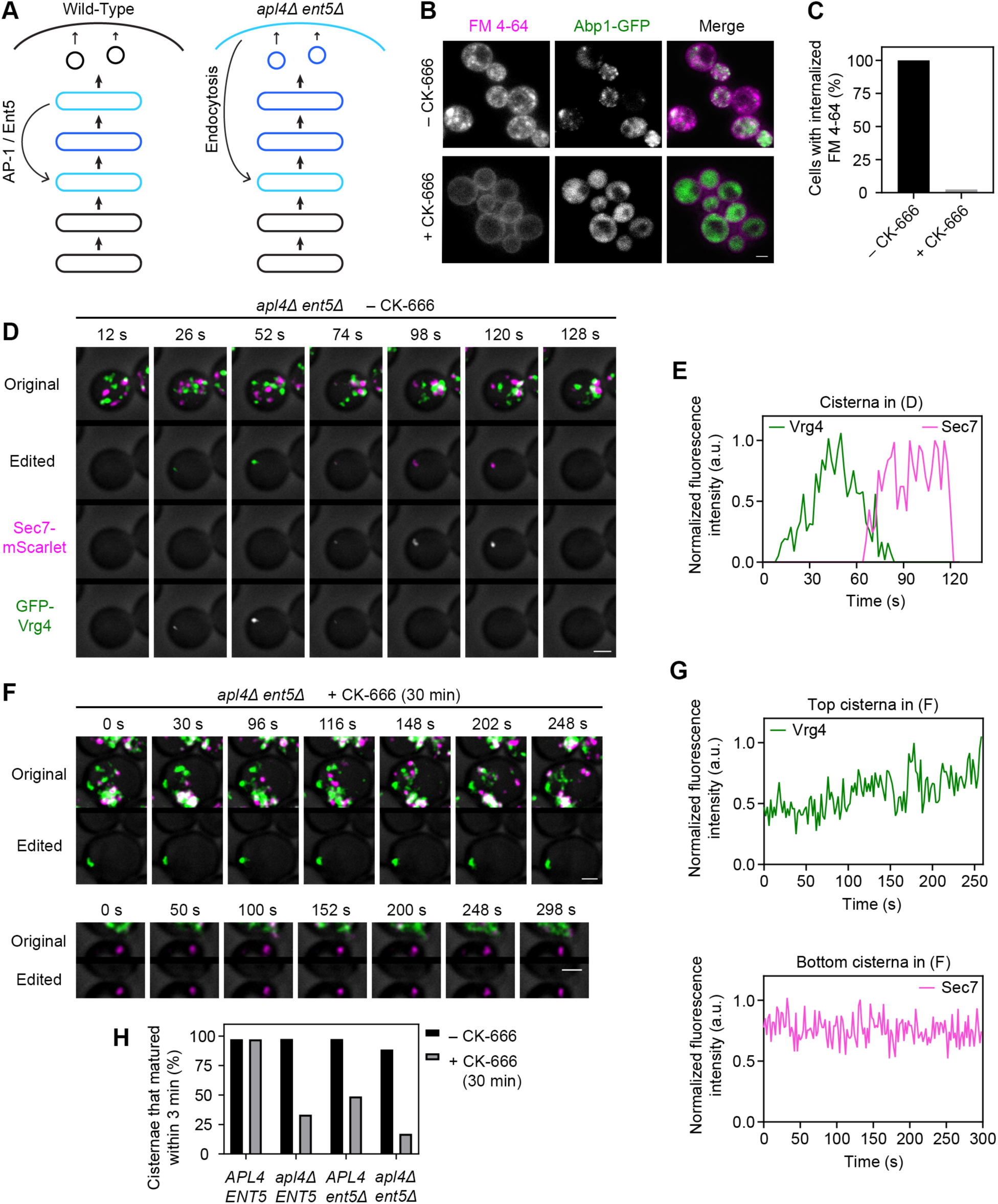
Inhibition of Endocytosis Blocks Golgi Maturation in Cells Lacking AP-1/Ent5. See also Video S2. (A) Diagram of the experimental rationale. In wild-type cells, a set of transmembrane TGN proteins, indicated in blue, recycle with the aid of AP-1/Ent5 from terminally maturing TGN cisternae to nascent TGN cisternae. In *apl4Δ ent5Δ* cells, which lack AP-1/Ent5, those TGN proteins presumably reach the plasma membrane in secretory vesicles and then undergo endocytic recycling to nascent TGN cisternae. Therefore, inhibition of endocytosis in an *apl4Δ ent5Δ* strain should trap transmembrane TGN proteins at the plasma membrane and perturb Golgi function. (B) Inhibition of endocytosis with CK-666. Cells expressing Abp1-GFP were mock treated or incubated with CK-666 for 15 min, then incubated for 5 min with FM 4-64FX, and then imaged by confocal microscopy. Shown are average projected Z-stacks. Individual fluorescence channels are shown in grayscale with merged images on the right. Scale bar, 2 µm. (C) Quantification of the analysis in (B). Projected images were manually scored for the presence of internalized dye. At least 40 cells were analyzed for each sample. (D) A typical Golgi maturation event in an untreated cell lacking AP-1 and Ent5. *apl4Δ ent5Δ* cells expressing the early Golgi marker GFP-Vrg4 and the TGN marker Sec7-mScarlet were grown to mid-log phase and imaged by 4D confocal microscopy. Shown are average projected Z-stacks at the indicated time points from Part 1 of Video S2. The upper row shows the complete projections, the second row shows edited projections that include only the cisterna being tracked, and the subsequent rows show the individual fluorescence channels from the edited projections. Scale bar, 2 µm. (E) Quantification of the fluorescence signals from the cisterna analyzed in (D). (F) Persistence of early or TGN markers in cisternae of *apl4Δ ent5Δ* cells after CK-666 treatment. Cells grown to mid-log phase were treated with CK-666 for 30 min prior to imaging as in (D). Shown are average projected Z-stacks at the indicated time points for separate cisternae from Parts 2 and 3 of Video S2. For each analyzed cisterna, the upper row shows the complete projections, and the lower row shows edited projections that include only the cisterna being tracked. Scale bars, 2 µm. (G) Quantification of the fluorescence signals from the cisternae analyzed in (F). (H) Quantification of Golgi maturation events in the absence or presence of CK-666. Cells expressing GFP-Vrg4 and Sec7-mScarlet in the indicated genetic backgrounds were either mock treated or treated with CK-666 for 30 min, and were imaged as in (D). Individual cisternae were scored according to whether or not they matured within 3 min. At least 36 cisternae from at least 29 cells were analyzed for each strain.

Our readout for Golgi function was cisternal maturation, as indicated by conversion of early Golgi cisternae labeled with the GDP-mannose transporter Vrg4 to TGN cisternae labeled with Sec7 (Losev et al., 2006). Normal maturation events were observed in untreated *apl4Δ ent5Δ* cells and in wild-type cells treated with CK-666 (Figure 2D,E,H and Video S2). However, in *apl4Δ ent5Δ* cells treated with CK-666, cisternal maturation was blocked. Various cisternae showed abnormal persistence of either Vrg4 or Sec7 (Figure 2F,G and Video S2), or more complex patterns that seemed to reflect hybrid or clustered cisternae (unpublished data). These effects are summarized in Figure 2H, which quantifies the percentage of cisternae that underwent maturation during a 3-min period. Removal of either AP-1 or Ent5 inhibited maturation in cells treated with CK-666, and this inhibition was nearly complete in cells lacking both adaptors. We conclude that components needed for Golgi operation recycle intracellularly in wild-type cells but recycle via the plasma membrane in *apl4Δ ent5Δ* cells.

### A set of transmembrane TGN proteins require AP-1/Ent5 for normal recycling

There is solid evidence that yeast AP-1 mediates recycling of the TGN-localized phospholipid translocase Drs2 (Liu et al., 2008). Based on our observation that AP-1-dependent recycling delivers material when a cisterna is beginning to acquire TGN characteristics (Casler et al., 2019), we predicted that Drs2 would arrive at a cisterna around the same time as Sec7. This prediction was confirmed by individual and averaged fluorescence traces, which revealed that Drs2 arrived shortly before Sec7 and then departed when AP-1 was present (Figure 3A-C and Video S3).

**Figure 3.**
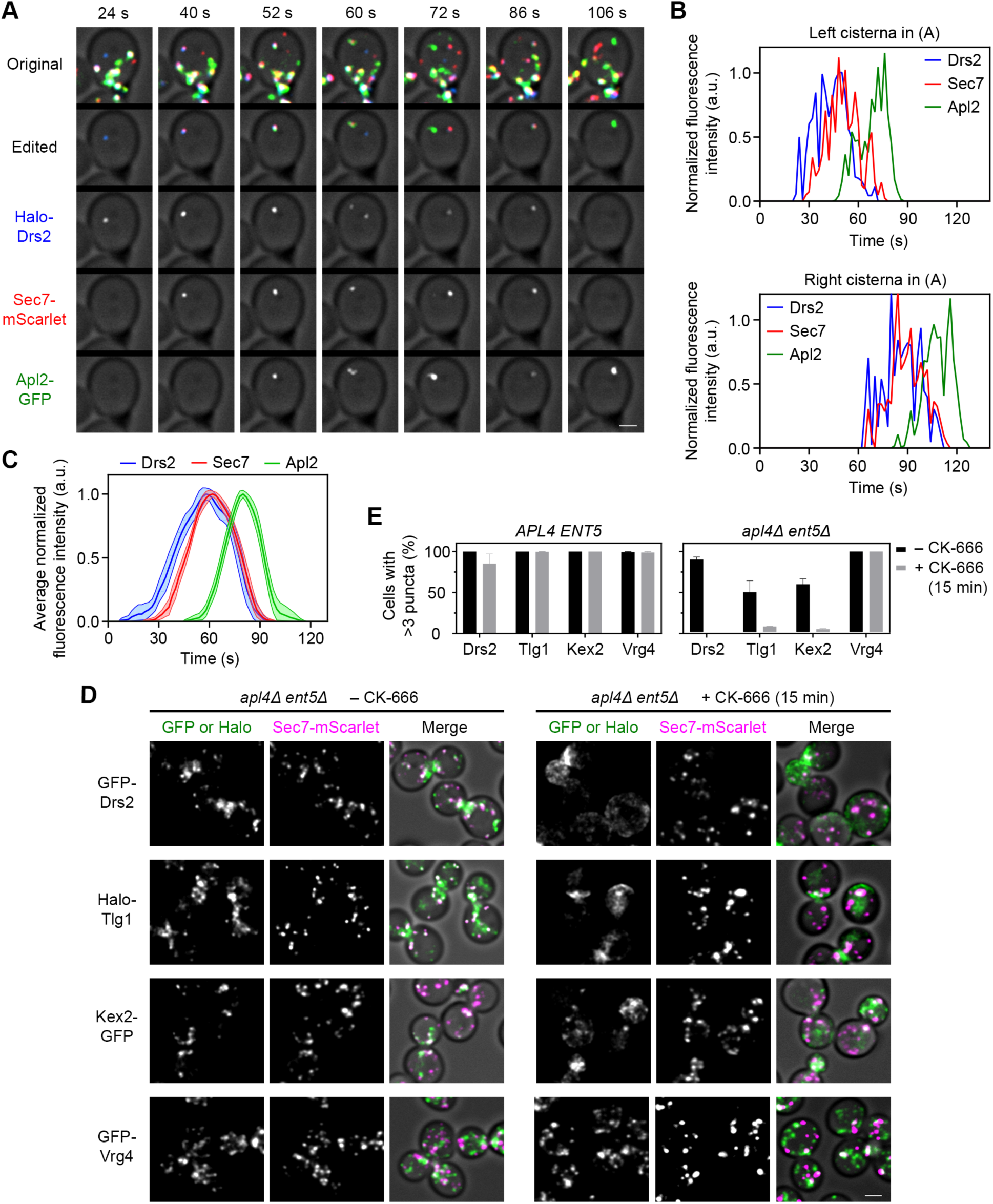
Drs2 and Other Transmembrane TGN Proteins Localize with the Aid of AP-1/Ent5. See also Figure S1, Figure S2, Figure S3, and Video S3. (A) Maturation kinetics of Drs2 compared to Sec7 and AP-1. A strain expressing the TGN marker Sec7-mScarlet, the AP-1 subunit Apl2-GFP, and HaloTag-Drs2 was grown to mid-log phase, labeled with JFX dye, and imaged by 4D confocal microscopy. Shown are average projected Z-stacks at the indicated time points from Video S3. The upper row shows the complete projections, the second row shows edited projections that include only the cisternae being tracked, and the subsequent rows show the individual fluorescence channels from the edited projections. Two events are shown. Scale bar, 2 µm. (B) Quantification of tagged Golgi proteins during typical maturation events. Depicted are the normalized fluorescence intensities in arbitrary units (a.u.) for the cisternae tracked in (A). (C) Smoothed and averaged traces showing the relative kinetic signatures of Drs2, Sec7, and Apl2. Data were obtained for 17 representative cisternae. (D) Mislocalization of transmembrane TGN proteins in *apl4Δ ent5Δ* cells after CK-666 treatment. *apl4Δ ent5Δ* cells expressing Sec7-mScarlet and the indicated HaloTag- or GFP-tagged Golgi protein were grown to mid-log phase, and then imaged by confocal microscopy 15 min after mock treatment or treatment with CK-666. Shown are average projected Z-stacks. Individual fluorescence channels are shown in grayscale with merged images on the right. Scale bar, 2 µm. (E) Quantification of the effects of the procedure in (D) for *APL4 ENT5* and *apl4Δ ent5Δ* cells. Average projected images were manually scored for the presence of 3 or more punctate spots in the GFP or HaloTag channel. Each bar represents an average of two biological replicates in which at least 40 cells were scored per condition. Error bars represent SEM.

As a functional test of whether AP-1/Ent5 mediates recycling of Drs2, we confirmed a report that in cells lacking AP-1, Drs2 accumulated at the plasma membrane when endocytosis was blocked (Liu et al., 2008). Our analysis involved treatment of wild-type or mutant cells with CK-666 for 15 min. When wild-type cells were treated with CK-666, Drs2 retained its punctate TGN localization (Figure S1A), but when *apl4Δ* or *apl4Δ ent5Δ* cells were treated with CK-666, Drs2 redistributed to the plasma membrane (Figure S2A and Figure 3D). We found that for a labeled Golgi protein such as Drs2, partial or full redistribution out of Golgi compartments could be quantified by counting the percentage of cells in which the labeled protein was visible in more than three punctate structures (Figure 3E). As a control, Vrg4 recycles in the early Golgi with the aid of COPI (Abe et al., 2004; Papanikou et al., 2015), and its distribution in either wild-type or *apl4Δ ent5Δ* cells was unaffected by CK-666 (Figure S1B and Figure 3D,E). Thus, CK-666 treatment of *apl4Δ ent5Δ* cells appears to be a suitable test of whether a transmembrane Golgi protein recycles with the aid of AP-1/Ent5.

We applied this test to several transmembrane TGN proteins that were candidates for AP-1/Ent5-mediated recycling based on their substantial colocalization with Sec7 (Figure S1A). A functional study implicated AP-1 in TGN localization of the SNARE protein Tlg1 (Valdivia et al., 2002), and when *apl4Δ ent5Δ* cells were treated with CK-666, the punctate intracellular distribution of Tlg1 was largely abolished (Figure 3D,E). The next candidate was the processing protease Kex2 (Fuller et al., 1988). It has long been thought that Kex2 recycles via PVE compartments, but the evidence is ambiguous (see Discussion), and we have proposed instead that Kex2 follows an intra-Golgi recycling pathway (Papanikou et al., 2015). When *apl4Δ ent5Δ* cells were treated with CK-666, the punctate intracellular distribution of Kex2 was largely abolished (Figure 3D,E), suggesting that AP-1/Ent5-dependent recycling is the primary localization mechanism for Kex2. Similar results were obtained with two other transmembrane Golgi proteins: Ste13, which is a dipeptidyl aminopeptidase that acts downstream of Kex2 (Fuller et al., 1988), and Stv1, which is a component of the TGN-localized proton-pumping ATPase (Manolson et al., 1994; Finnigan et al., 2012) (Figure S3A). In wild-type cells, as expected, CK-666 had no effect on the punctate TGN localizations of Tlg1, Kex2, Ste13, or Stv1 (Figure S1A). Further control experiments confirmed that the early Golgi glycosylation enzymes Anp1 and Mnn9 (Jungmann and Munro, 1998) resembled Vrg4 in being unaffected by CK-666 in either wild-type or *apl4Δ ent5Δ* cells (Figure S1B and Figure S3B). Prior to CK-666 treatment, *apl4Δ ent5Δ* cells showed reduced TGN labeling for Drs2, Tlg1, Kex2, Ste13, and Stv1—presumably because some of the protein molecules were recycling between the TGN and plasma membrane at steady-state—whereas no such effect was seen for Vrg4, Anp1, or Mnn9 (Figure S3A,B and unpublished data). Thus, AP-1/Ent5 specifically mediates recycling of a set of transmembrane TGN proteins.

These TGN proteins did exhibit some differences in their responses. After CK-666 treatment, Drs2 showed an exceptionally clean redistribution to the plasma membrane in an *apl4Δ ent5Δ* strain (Figure 3D), and Drs2 uniquely showed extensive redistribution to the plasma membrane in either an *apl4Δ* strain or an *ent5Δ* strain (Figure S2A,B and Figure S3A). We speculate that endocytosis of Drs2 is relatively inefficient, so any increased leakage of Drs2 into secretory vesicles leads to kinetic trapping at the plasma membrane. Other TGN proteins responded to CK-666 treatment of *apl4Δ ent5Δ* cells by combining plasma membrane labeling with accumulation at sites of polarized secretion (Figure 3D and unpublished data). In an *ent5Δ* strain but not in an *apl4Δ* strain, Tlg1 showed decreased TGN localization in untreated cells (Hung and Duncan, 2016) and loss of TGN localization in CK-666-treated cells (Figure S2A,B and Figure S3A), suggesting that Ent5 is especially important for Tlg1 localization. Despite these variations, the basic result is that removal of AP-1/Ent5 alters the trafficking of multiple transmembrane TGN proteins.

### AP-1/Ent5-dependent TGN proteins show similar kinetic signatures

We compared the kinetic signatures of the AP-1/Ent5-dependent Tlg1, Kex2, Ste13, and Stv1 proteins to that of Drs2 during Golgi maturation (Figure 4A-F and Figure S4A-F and Video S4). All four TGN proteins resembled Drs2 in their kinetic signatures. Drs2 tended to persist about 5-15 s longer than the other TGN proteins, perhaps because it is packaged less efficiently into AP-1/Ent5 carriers, but those differences were minor. By contrast, the AP-1/Ent5-independent early Golgi proteins Vrg4, Anp1, and Mnn9 arrived and departed about 60 s earlier than Drs2 (Figure 4G-I and Figure S5 and Video S5). These data support the assignment of Drs2, Tlg1, Kex2, Ste13, and Stv1 to a kinetic class that reflects their recycling pathway.

**Figure 4.**
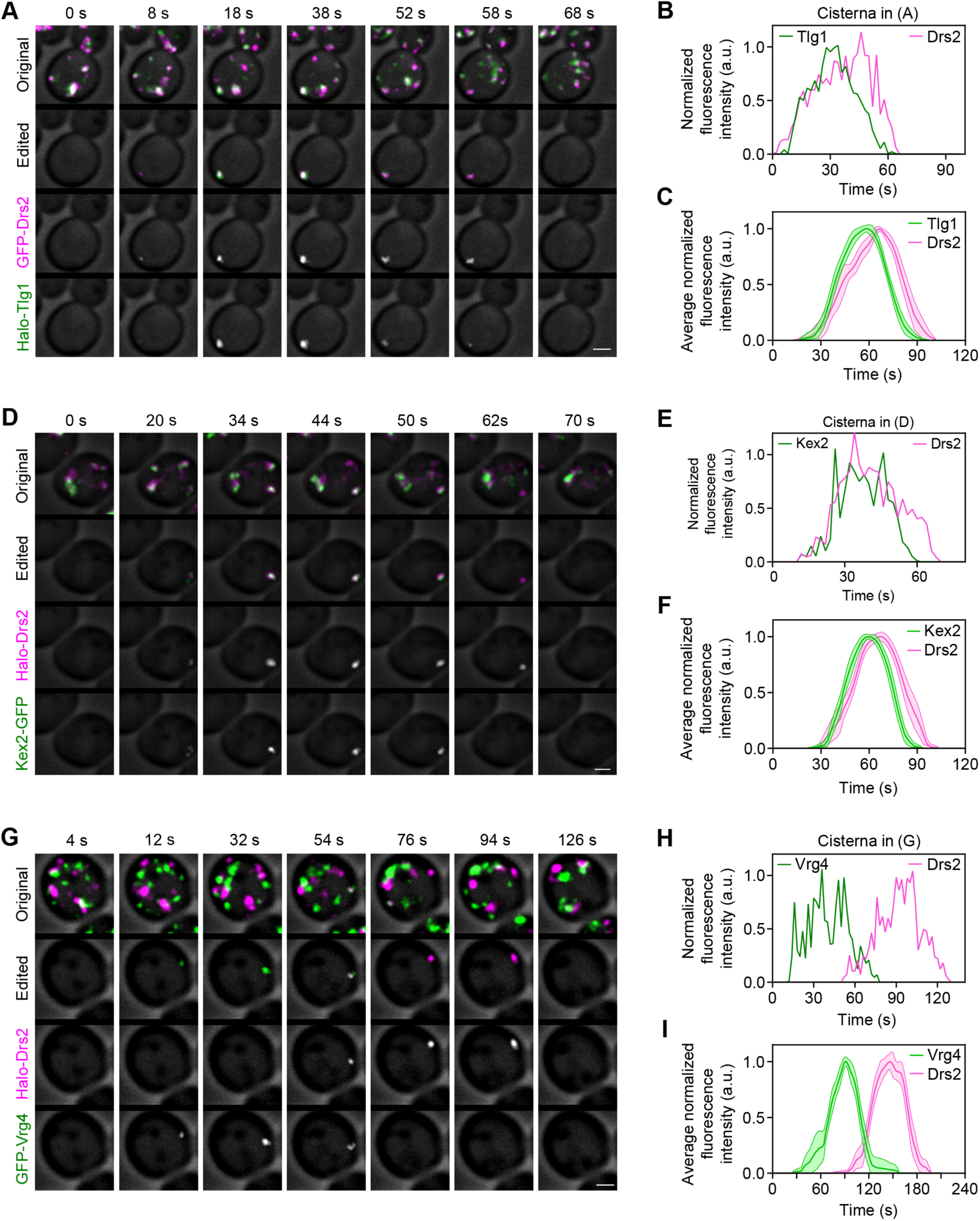
Other Transmembrane TGN Proteins Resemble Drs2 in Their Kinetic Signatures. See also Figure S4, Figure S5, Video S4, and Video S5. (A) Maturation kinetics of Drs2 compared to Tlg1. A strain expressing GFP-Drs2 and the SNARE HaloTag-Tlg1 was grown to mid-log phase, labeled with JFX dye, and imaged by 4D confocal microscopy. Shown are average projected Z-stacks at the indicated time points from Part 1 of Video S4. The upper row shows the complete projections, the second row shows edited projections that include only the cisterna being tracked, and the subsequent rows show the individual fluorescence channels from the edited projections. Scale bar, 2 µm. (B) Quantification of tagged Golgi proteins during a typical maturation event. Depicted are the normalized fluorescence intensities in arbitrary units (a.u.) for the cisterna tracked in (A). (C) Smoothed and averaged traces showing the relative kinetic signatures of Drs2 and Tlg1. Data were obtained for 10 representative cisternae. (D) Maturation kinetics of Drs2 compared to Kex2. The experiment was performed as in (A), except with a strain expressing HaloTag-Drs2 and the processing protease Kex2-GFP. Shown are average projected Z-stacks at the indicated time points from Part 2 of Video S4. Scale bar, 2 µm. (E) Quantification of tagged Golgi proteins during a typical maturation event. Depicted are the normalized fluorescence intensities in arbitrary units (a.u.) for the cisterna tracked in (D). (F) Smoothed and averaged traces showing the relative kinetic signatures of Drs2 and Kex2. Data were obtained for 13 representative cisternae. (G) Maturation kinetics of Drs2 compared to Vrg4. The experiment was performed as in (A), except with a strain expressing HaloTag-Drs2 and GFP-Vrg4. Shown are average projected Z-stacks at the indicated time points from Part 1 of Video S5. Scale bar, 2 µm. (H) Quantification of tagged Golgi proteins during a typical maturation event. Depicted are the normalized fluorescence intensities in arbitrary units (a.u.) for the cisterna tracked in (G). (I) Smoothed and averaged traces showing the relative kinetic signatures of Drs2 and Vrg4. Data were obtained for 12 representative cisternae.

### TGN proteins that recycle via PVE compartments depart much sooner than Drs2 and localize independently of AP-1/Ent5

Some transmembrane TGN proteins, such as the vacuolar hydrolase receptor Vps10, travel to PVE compartments and then recycle to the Golgi (Marcusson et al., 1994; Cooper and Stevens, 1996). Vps10 is packaged into Golgi-derived vesicles with the aid of the GGA clathrin adaptors (Black and Pelham, 2000; Dell’Angelica et al., 2000; Hirst et al., 2000; Zhdankina et al., 2001), which arrive before AP-1 and Ent5 (Daboussi et al., 2012; Casler and Glick, 2020). We showed recently that a Vps10-dependent biosynthetic cargo begins to exit the Golgi immediately after a cisterna acquires TGN characteristics (Casler and Glick, 2020). Based on these considerations, Vps10 would be expected to depart from TGN cisternae significantly before Drs2. Indeed, the kinetic signatures of Vps10 and Drs2 were quite different (Figure 5A-C and Video S6). Although Vps10 arrived at about the same time as Drs2, it accumulated more rapidly, and then began to depart while Drs2 levels were still increasing. As a second example of a TGN protein that likely recycles via PVE compartments, we examined the Na^+^/H^+^ exchanger Nhx1, which resembles Vps10 in showing dual localization to TGN and PVE structures (Kojima et al., 2012; Chi et al., 2014; Papanikou et al., 2015). The kinetic signature of Nhx1 was similar to that of Vps10 and clearly distinct from that of Drs2 (Figure 5D-F and Video S6). Thus, Vps10 and Nhx1 can be assigned to a second kinetic class of transmembrane TGN proteins.

**Figure 5.**
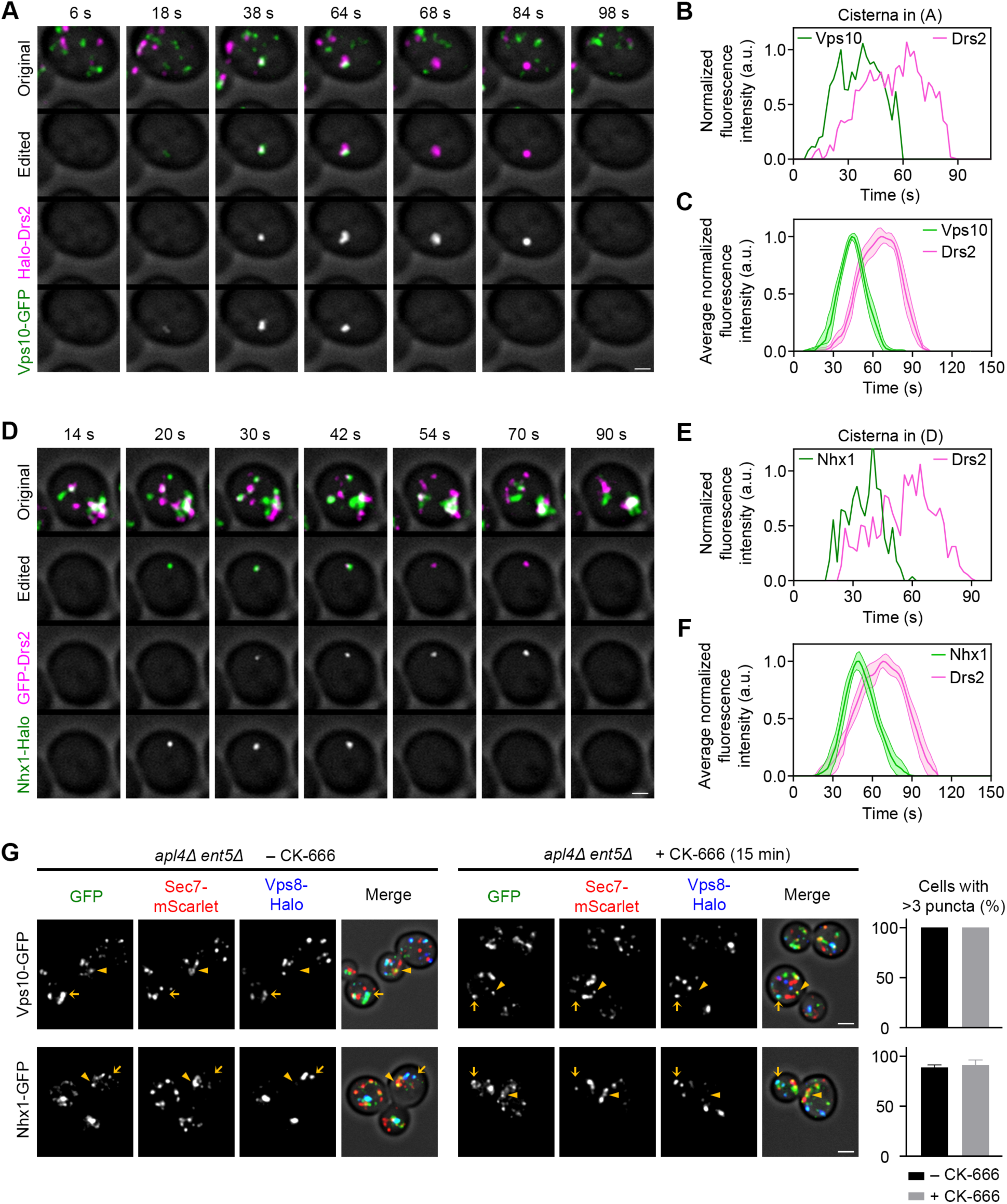
A Separate Class of Transmembrane TGN Proteins Recycle via PVE Compartments. See also Video S6. (A) Maturation kinetics of Drs2 compared to Vps10. A strain expressing HaloTag-Drs2 and the vacuolar hydrolase receptor Vps10-GFP was grown to mid-log phase, labeled with JFX dye, and imaged by 4D confocal microscopy. Shown are average projected Z-stacks at the indicated time points from Part 1 of Video S6. The upper row shows the complete projections, the second row shows edited projections that include only the cisterna being tracked, and the subsequent rows show the individual fluorescence channels from the edited projections. Scale bar, 2 µm. (B) Quantification of tagged Golgi proteins during a typical maturation event. Depicted are the normalized fluorescence intensities in arbitrary units (a.u.) for the cisterna tracked in (A). (C) Smoothed and averaged traces showing the relative kinetic signatures of Drs2 and Vps10. Data were obtained for 15 representative cisternae. (D) Maturation kinetics of Drs2 compared to Nhx1. The experiment was performed as in (A), except with a strain expressing GFP-Drs2 and the Na^+^/H^+^ exchanger Nhx1-HaloTag. Shown are average projected Z-stacks at the indicated time points from Part 2 of Video S6. Scale bar, 2 µm. (E) Quantification of tagged Golgi proteins during a typical maturation event. Depicted are the normalized fluorescence intensities in arbitrary units (a.u.) for the cisterna tracked in (D). (F) Smoothed and averaged traces showing the relative kinetic signatures of Drs2 and Nhx1. Data were obtained for 8 representative cisternae. (G) Dual localization of Vps10 and Nhx1 before and after treatment with CK-666. *apl4Δ ent5Δ* strains expressing the TGN marker Sec7-mScarlet, the PVE compartment marker Vps8-HaloTag, and either Vps10-GFP or Nhx1-GFP were grown to mid-log phase, labeled with JFX dye, and imaged by 4D confocal microscopy 15 min after mock treatment or treatment with CK-666. Shown are average projected Z-stacks. Individual fluorescence channels are shown in grayscale with merged images on the right. Arrowheads mark examples of Sec7-labeled TGN structures that contain Vps10 or Nhx1, and arrows mark examples of Vps8-labeled PVE compartments that contain Vps10 or Nhx1. Scale bar, 2 µm. At the right, the same *apl4Δ ent5Δ* strains were treated with CK-666 and manually scored for the presence of 3 or more punctate spots in the GFP channel, as in Figure 3E.

Because Vps10 exits the Golgi in GGA-dependent carriers, its intracellular distribution should be unaffected by removal of AP-1/Ent5. Indeed, when *apl4Δ ent5Δ* cells were treated with CK-666, Vps10 showed no loss of punctate intracellular localization (Figure 5G,H). Similar results were obtained for Nhx1 (Figure 5G,H). The localization of Vps10 and Nhx1 to both TGN and PVE structures was similar in wild-type and *apl4Δ ent5Δ* cells (Figure 5G and unpublished data). We conclude that Vps10 and Nhx1 recycle in a manner independent of AP-1/Ent5. These findings bolster the argument that the kinetic signatures of Golgi transmembrane proteins reflect their recycling pathways.

### A class of AP-1/Ent5-dependent Golgi proteins have intermediate residence times

We were curious about the transmembrane protein Sys1, which initiates the process of recruiting the Arl1 GTPase (Behnia et al., 2004; Setty et al., 2004). Our original analysis suggested that Sys1 and Sec7 had similar kinetic signatures (Losev et al., 2006), but for that experiment Sys1 was overexpressed, and later studies from the Nakano group reported that Sys1 arrives and departs well before Sec7 (Ishii et al., 2016; Kurokawa et al., 2019; Tojima et al., 2019). Figure 6A-C and Video S7 confirm the distinct kinetic signatures of Sys1 and Sec7, and also provide a comparison with Drs2. On average, Sys1 arrived and departed about 15-20 s before Drs2. Sys1 therefore defines an additional kinetic class of transmembrane Golgi proteins.

**Figure 6.**
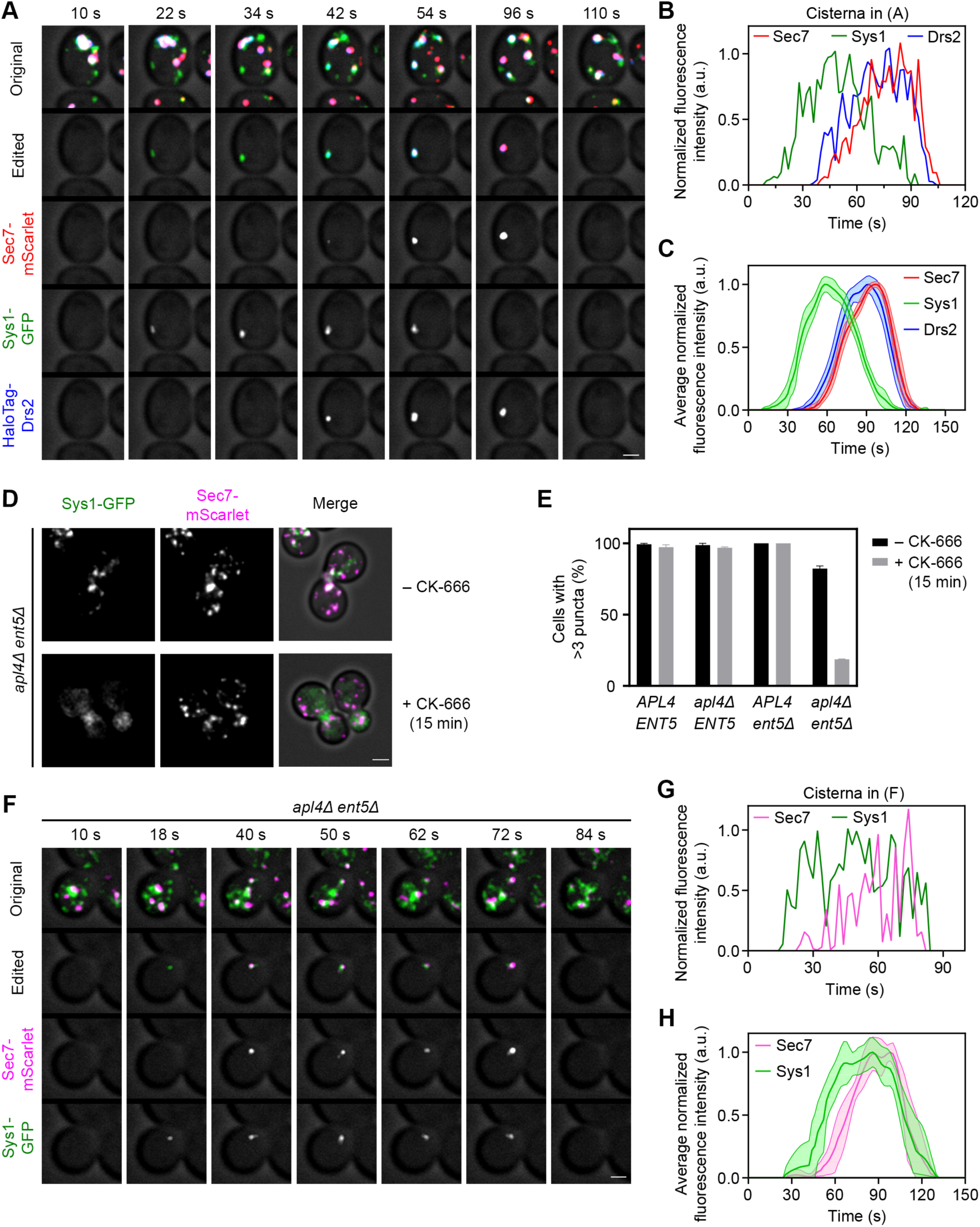
Sys1 Localizes with the Aid of AP-1/Ent5 But Has an Intermediate Kinetic Signature. See also Video S7. (A) Maturation kinetics of Sys1 compared to Sec7 and Drs2. A strain expressing Sec7-mScarlet, HaloTag-Drs2, and the transmembrane Golgi protein Sys1-GFP was grown to mid-log phase, labeled with JFX dye, and imaged by 4D confocal microscopy. Shown are average projected Z-stacks at the indicated time points from Part 1 of Video S7. The upper row shows the complete projections, the second row shows edited projections that include only the cisterna being tracked, and the subsequent rows show the individual fluorescence channels from the edited projections. Scale bar, 2 µm. (B) Quantification of tagged Golgi proteins during typical maturation events. Depicted are the normalized fluorescence intensities in arbitrary units (a.u.) for the cisterna tracked in (A). (C) Smoothed and averaged traces showing the relative kinetic signatures of Sec7, Sys1, and Drs2. Data were obtained for 10 representative cisternae. (D) Mislocalization of Sys1 in *apl4Δ ent5Δ* cells after CK-666 treatment. The experiment was performed as in Figure 3D. Scale bar, 2 µm. (E) Quantification of the effects in (D). The experiment was performed as in Figure 3E, except that *apl4Δ ENT5* and *APL4 ent5*Δ strains were also examined. (F) Maturation kinetics of Sys1 compared to Sec7 in an *apl4Δ ent5Δ* strain. The experiment was performed as in (A), except that HaloTag-Drs2 was not imaged because the signal was too weak. Shown are average projected Z-stacks at the indicated time points from Part 2 of Video S7. Scale bar, 2 µm. (G) Quantification of tagged Golgi proteins during typical maturation events in an *apl4Δ ent5Δ* strain. Depicted are the normalized fluorescence intensities in arbitrary units (a.u.) for the cisterna tracked in (F). (H) Smoothed and averaged traces showing the relative kinetic signatures of Sec7 and Sys1 in an *apl4Δ ent5Δ* strain. Data were obtained for 7 representative cisternae.

Unexpectedly, when *apl4Δ ent5Δ* cells were treated with CK-666, the punctate Golgi localization of Sys1 was lost (Figure 6D,E). This effect was seen only upon removal of both AP-1 and Ent5 (Figure 6E). The implication is that in the absence of AP-1/Ent5, Sys1 switches to the bypass recycling pathway of transit to the plasma membrane followed by endocytosis. We found previously that endocytic vesicles arrive at Golgi cisternae shortly before Sec7 (Day et al., 2018). Thus, in *apl4Δ ent5Δ* cells, Sys1 should arrive shortly before Sec7, and should persist during TGN maturation until being packaged into secretory vesicles. When we tested this prediction, the data were noisy because the Golgi signal for Sys1 was reduced in the absence of AP-1/Ent5, but there was a major shift that brought the Sys1 kinetic signature much closer to that of Sec7 (Figure 6F-H and Video S7). Quantification of the averaged data revealed that in wild-type cells, Sys1 arrived ∼25 s before Sec7 and departed ∼19 s before Sec7, whereas in *apl4Δ ent5Δ* cells, Sys1 arrived ∼7 s before Sec7 and departed at the same time as Sec7. A plausible interpretation is that AP-1/Ent5 mediates a second intra-Golgi recycling pathway that enables Sys1 to reside in cisternae during an intermediate phase of maturation.

Do other transmembrane Golgi proteins recycle in the same manner as Sys1? We examined Golgi proteins that have been described as having neither early nor TGN localizations. One candidate was Aur1, an inositol phosphorylceramide synthase (Nagiec et al., 1997; Levine et al., 2000). Aur1 reportedly shows partial colocalization with early and TGN markers (Levine et al., 2000). Another candidate was Rbd2, a putative rhomboid protease that also shows partial colocalization with early and TGN markers (Cortesio et al., 2015; Lastun et al., 2016). We found that both Aur1 and Rbd2 had kinetic signatures resembling that of Sys1 (Figure 7C-H and Video S8). Moreover, both proteins redistributed out of punctate Golgi structures when *apl4Δ ent5Δ* cells were treated with CK-666 (Figure 7A,B). The loss of Golgi localization was less extensive for Aur1 and Rbd2 (Figure 7B) than for the transmembrane TGN proteins examined earlier (Figure S3A), but the effect was clear. Therefore, AP-1/Ent5 is involved in recycling not only TGN proteins, but also a class of transmembrane proteins that would traditionally be said to reside in the medial/*trans* Golgi.

**Figure 7.**
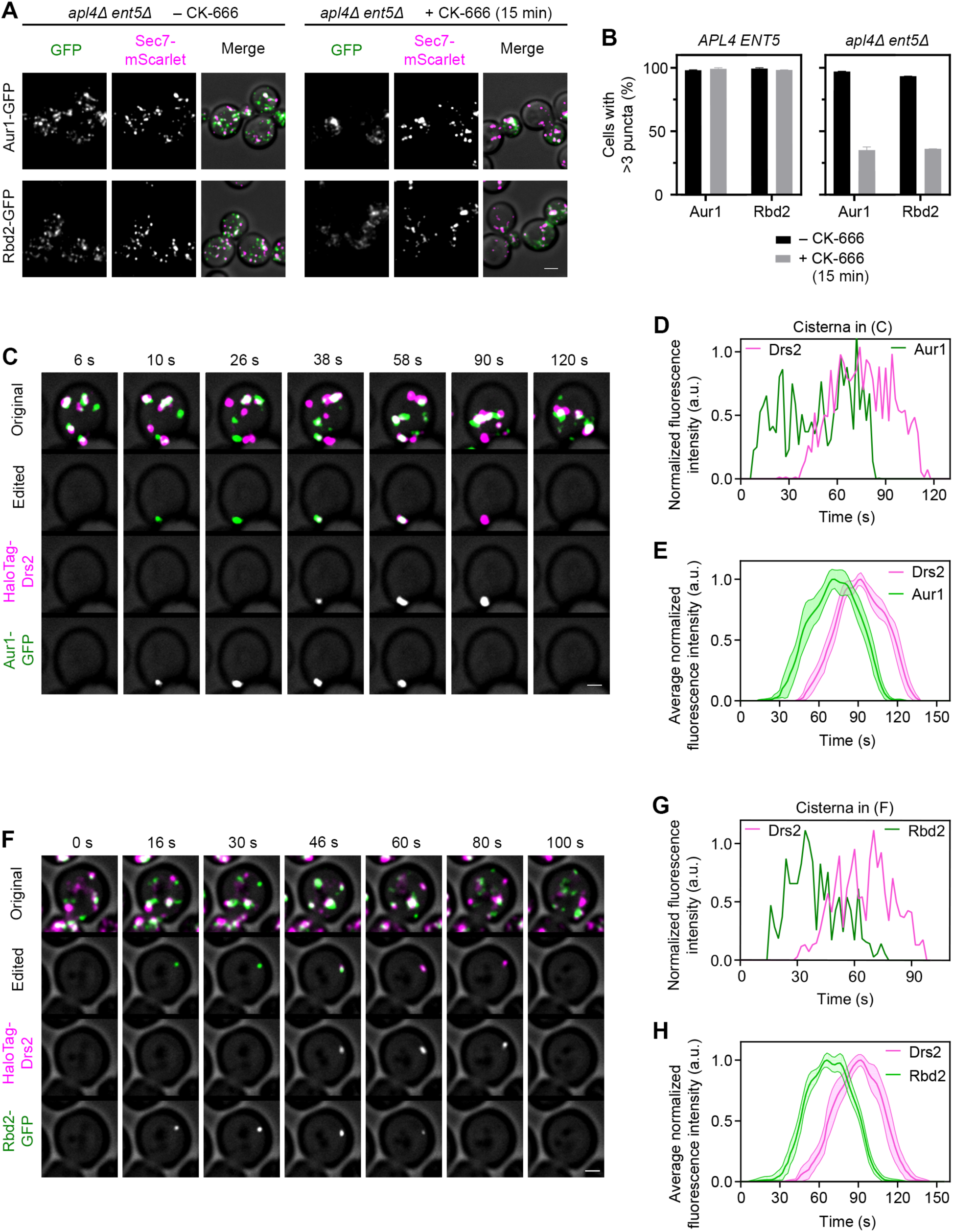
Other Transmembrane Golgi Proteins Resemble Sys1 in Their Kinetic Signatures. See also Video S8. (A) Mislocalization of the transmembrane Golgi proteins Aur1 and Rbd2 in *apl4Δ ent5Δ* cells after CK-666 treatment. The experiment was performed as in Figure 3D. Scale bar, 2 µm. (B) Quantification of the effects in (A). The experiment was performed as in Figure 3E. (C) Maturation kinetics of Drs2 compared to Aur1. A strain expressing HaloTag-Drs2 and Aur1-GFP was grown to mid-log phase, labeled with JFX dye, and imaged by 4D confocal microscopy. Shown are average projected Z-stacks at the indicated time points from Part 1 of Video S8. The upper row shows the complete projections, the second row shows edited projections that include only the cisterna being tracked, and the subsequent rows show the individual fluorescence channels from the edited projections. Scale bar, 2 µm. (D) Quantification of tagged Golgi proteins during typical maturation events. Depicted are the normalized fluorescence intensities in arbitrary units (a.u.) for the cisterna tracked in (C). (E) Smoothed and averaged traces showing the relative kinetic signatures of Drs2 and Aur1. Data were obtained for 10 representative cisternae. (F) Maturation kinetics of Drs2 compared to Rbd2. The experiment was performed as in (C), except with a strain expressing HaloTag-Drs2 and Rbd2-GFP. Shown are average projected Z-stacks at the indicated time points from Part 2 of Video S8. Scale bar, 2 µm. (G) Quantification of tagged Golgi proteins during typical maturation events. Depicted are the normalized fluorescence intensities in arbitrary units (a.u.) for the cisterna tracked in (F). (H) Smoothed and averaged traces showing the relative kinetic signatures of Drs2 and Rbd2. Data were obtained for 14 representative cisternae.

## Discussion

The polarized distribution of transmembrane proteins in the Golgi has long been mysterious. Previously postulated mechanisms include trapping of Golgi proteins in cisternae by association with other proteins of the same type (Nilsson et al., 1993), or partitioning of Golgi proteins into different regions of the organelle based on transmembrane domain properties (Banfield, 2011; Welch and Munro, 2019), or competition between Golgi proteins for packaging into retrograde COPI vesicles (Glick et al., 1997). We have proposed that the key factor is recycling pathways—for a given transmembrane Golgi protein, the recycling pathway determines when the protein resides in a maturing cisterna (Pantazopoulou and Glick, 2019). Thus, each recycling pathway will define a class of transmembrane Golgi proteins. Proteins of different classes will show distinct kinetic signatures and distinct patterns of concentration in the cisternae (Figure S6A).

To test this idea with yeast cells, we need to know the recycling pathways that operate at the Golgi. One pathway, not examined here, is COPI-dependent transport from the early Golgi to the ER followed by recycling to the early Golgi (Gaynor et al., 1998; Barlowe and Miller, 2013). A second pathway is COPI-dependent recycling within the early Golgi of proteins such as Vrg4 (Papanikou et al., 2015) (Figure S6B). A third pathway involves AP-1. Functional studies originally suggested that yeast AP-1 recycles transmembrane proteins from early endosomes to the TGN (Valdivia et al., 2002; Foote and Nothwehr, 2006; Liu et al., 2008; Spang, 2015). However, yeast early endosomes are identical to the TGN, and AP-1 localizes exclusively to maturing TGN cisternae, implying that AP-1 mediates intra-Golgi recycling of TGN proteins (Day et al., 2018). An example of an AP-1-dependent TGN protein is Drs2 (Liu et al., 2008). We show here that Drs2 departs from maturing cisternae after AP-1 appears, and then arrives at younger cisternae shortly before the early-to-TGN transition. The combined results imply that AP-1 mediates intra-Golgi recycling of Drs2 downstream of COPI (Papanikou et al., 2015; Day et al., 2018; Casler et al., 2019).

A well-characterized TGN protein is Kex2 (Fuller et al., 1988). Our earlier video microscopy studies (Papanikou et al., 2015) were extended here by showing that the kinetic signature of Kex2 resembles that of Drs2. To test whether Kex2 relies on AP-1 for its localization, we built on a previously described strategy (Valdivia et al., 2002; Liu et al., 2008). The rationale is that if AP-1 mediates recycling of a Golgi protein, then removal of AP-1 should divert that Golgi protein to the cell surface, where it can be trapped by inhibiting endocytosis. We optimized a procedure for specifically inhibiting endocytosis by using CK-666 to block formation of branched actin (Hetrick et al., 2013; Burke et al., 2014; Antkowiak et al., 2019). In cells lacking AP-1, CK-666 trapped Drs2 at the plasma membrane, consistent with a published report (Liu et al., 2008). However, we did not see a similar effect for Kex2 or other transmembrane proteins that were candidates for recycling within the TGN. One possible explanation was that in addition to AP-1, another recycling factor might operate at the TGN.

The epsin-related adaptors Ent3, Ent4, and Ent5 localize to the yeast TGN (Myers and Payne, 2013). Little is known about Ent4 (Deng et al., 2009). Ent3 appears to act early in TGN maturation together with GGAs (Costaguta et al., 2006; Copic et al., 2007; Daboussi et al., 2012). Ent5 also interacts with GGAs, but the primary role of Ent5 seems to be later in TGN maturation, when it acts together with AP-1 (Costaguta et al., 2006; Copic et al., 2007; Daboussi et al., 2012; Hung and Duncan, 2016). We therefore tested whether the AP-1/Ent5 pair mediates intra-Golgi recycling. Indeed, in cells lacking AP-1/Ent5, CK-666 caused redistribution of Kex2 and several other TGN proteins out of Golgi structures. No such effect was seen for early Golgi proteins or for Golgi proteins that recycle from PVE compartments. The implication is that in the mutant cells, proteins that would normally recycle with the aid of AP-1/Ent5 travel instead to the plasma membrane and then return to the Golgi by endocytosis. This concept explains why removal of AP-1/Ent5 confers only a mild phenotype, and it provides a convenient way to test whether a given transmembrane Golgi protein undergoes AP-1/Ent5-dependent recycling.

We predicted that the AP-1/Ent5-dependent TGN proteins would all have similar kinetic signatures. This prediction was largely confirmed. Drs2 tended to depart somewhat later during maturation than other TGN proteins, suggesting that packaging into AP-1/Ent5 carriers either is regulated, or is more efficient for some TGN proteins than others. But overall, the AP-1/Ent5-dependent class of TGN proteins showed only small differences in their kinetic signatures (Figure S6B), consistent with the idea that the recycling pathway determines when a transmembrane Golgi protein is present in maturing cisternae.

Further support for this concept came from examining TGN proteins that follow a different recycling pathway. Vps10 undergoes GGA-dependent traffic from the TGN to PVE compartments followed by recycling to the Golgi (Marcusson et al., 1994; Cooper and Stevens, 1996), so Vps10 is found both in the TGN and in PVE compartments (Chi et al., 2014; Papanikou et al., 2015). As expected, Vps10 localization is unperturbed when cells lacking AP-1/Ent5 are treated with CK-666. Vps10 resembles a previously characterized Vps10-dependent cargo (Casler and Glick, 2020) in departing to PVE compartments soon after the early-to-TGN transition. Because Vps10 arrives at cisternae just before the early-to-TGN transition, this protein has a relatively short residence time during cisternal maturation. The kinetic signature of Vps10 overlaps with that of Drs2 (orange versus blue curves in Figure S6A), but Vps10 departs earlier than Drs2 because GGAs act before AP-1/Ent5 (Daboussi et al., 2012; Casler and Glick, 2020). Similar results were obtained for Nhx1, which is also believed to cycle between the TGN and PVE compartments (Kojima et al., 2012). Therefore, we can differentiate between two classes of TGN proteins based on their recycling pathways and kinetic signatures (Figure S6B).

The assignment of Kex2 to a different class of TGN proteins than Vps10 was not obvious because for many years, Kex2 has been assumed to recycle from PVE compartments. As evidence for that view, when mutations were used to block PVE-to-Golgi recycling, Kex2 accumulated in PVE compartments and the vacuole (Wilcox et al., 1992; Voos and Stevens, 1998). But those results are potentially misleading because if Kex2 visits PVE compartments only occasionally, a block in recycling will still trap most of the Kex2 molecules in PVE compartments. Unlike Vps10, Kex2 has no known function in PVE compartments and is present at very low concentrations in PVE compartments, suggesting that the main pathway for Kex2 localization might be recycling within the TGN (Papanikou et al., 2015; Day et al., 2018). Similar arguments hold for Ste13, a TGN protein that acts after Kex2 (Fuller et al., 1988). Ste13 has also been assumed to recycle from PVE compartments, and indeed, the signals and mechanisms for retrieving Ste13 from PVE compartments have been extensively studied (Voos and Stevens, 1998; Nothwehr et al., 2000; Harrison et al., 2014; Ma and Burd, 2020). Yet our analysis classifies Ste13 together with Kex2 as an AP-1/Ent5-dependent TGN protein. An elegant pair of earlier reports showed that the transit of Ste13 to PVE compartments is slow with a half-time of ∼60 min, and that a fast-acting AP-1-dependent process keeps Ste13 in the TGN (Bryant and Stevens, 1997; Foote and Nothwehr, 2006). We suggest that proteins such as Kex2 and Ste13 normally recycle within the TGN by an AP-1/Ent5-dependent pathway, but they occasionally escape to PVE compartments, where a salvage pathway returns them to the TGN.

The hypothesis that COPI recycles early Golgi proteins while AP-1/Ent5 recycles many TGN proteins does not account for Sys1, which was shown to have an intermediate kinetic signature (Ishii et al., 2016; Kurokawa et al., 2019; Tojima et al., 2019). We verified that Sys1 arrives and departs earlier than Drs2 and other TGN proteins, and we observed the same type of intermediate kinetic signature for two other transmembrane Golgi proteins (Figure S6B). Remarkably, in cells lacking AP-1/Ent5, CK-666 caused redistribution of proteins such as Sys1 out of Golgi structures. We infer that AP-1/Ent5 participates in the recycling of two classes of Golgi proteins by two kinetically distinct pathways.

A simple explanation would be that one class of Golgi proteins is recycled by AP-1 and the other by Ent5, but for most of the Golgi proteins tested, CK-666 had little effect in cells singly lacking either AP-1 or Ent5. The relationship between AP-1 and Ent5 is complex and ambiguous. These two adaptors physically interact and yet can function independently (Costaguta et al., 2006; Copic et al., 2007). In our hands, Ent5 departs from the TGN sooner than AP-1, suggesting that AP-1 and Ent5 are not merely components of the same carriers. We suggest that either AP-1 or Ent5 or the AP-1/Ent5 pair can recycle transmembrane TGN proteins plus an intermediate class of transmembrane Golgi proteins.

Do these two kinetic classes of transmembrane Golgi proteins actually define two separate AP-1/Ent5-mediated recycling pathways? This idea has a precedent with COPI, which is a single vesicle-forming complex that apparently mediates both Golgi-to-ER and intra-Golgi recycling (Popoff et al., 2011). An alternative possibility is that various Golgi proteins access the same AP-1/Ent5-dependent recycling pathway at different times. However, the kinetic signatures of the AP-1/Ent5-dependent proteins seem to define two kinetic classes rather than a continuum, suggesting the existence of two separate pathways. Perhaps the transmembrane Golgi proteins that follow these pathways are not passive passengers, but are actively involved in harnessing the AP-1/Ent5 machinery to create two types of carriers. A caveat to this interpretation is the lack of evidence that specific Golgi proteins are packaged into carriers containing AP-1 and/or Ent5. Another concern is that proteins such as Sys1 begin to depart from a maturing cisterna when the levels of membrane-associated AP-1 and Ent5 are very low. This discrepancy can potentially be explained if early-arriving AP-1 and Ent5 molecules are rapidly packaged into carriers that bud from the cisternal membrane, in which case the activity of an adaptor would precede its accumulation on a cisterna. We saw such a phenomenon with a GGA-dependent vacuolar protein, which often began to depart from a maturing cisterna before a GGA adaptor showed significant accumulation (Casler and Glick, 2020). Yet another possibility is that the AP-1/Ent5 effects on proteins such as Sys1 are indirect. For example, loss of AP-1/Ent5 could inhibit a previously unknown GGA-mediated intra-Golgi recycling pathway. To clarify the mechanisms of the two AP-1/Ent5-dependent recycling pathways, we will seek to determine which vesicle tethers are involved in each pathway, and will then ectopically localize those tethers to capture the two classes of AP-1/Ent5-dependent carriers (Wong and Munro, 2014).

A longer-term project will be to determine why the cell opts to recycle particular transmembrane Golgi proteins by specific pathways. As an example, there may be a functional explanation for why Sys1 arrives before Drs2. Sys1 initiates the recruitment of Arl1, which in turn recruits the Imh1 tether (Panic et al., 2003; Setty et al., 2003), so does Imh1 then capture the carriers that recycle Drs2? Answers to such questions will shed light on the mechanisms that choreograph Golgi maturation.

Golgi trafficking components tend to be highly conserved. In mammalian cells, the data are consistent with AP-1-dependent retrograde traffic at the TGN (Hinners and Tooze, 2003; Hirst et al., 2012). Mammalian cells also contain a Golgi-localized epsin-related protein called epsinR that may function similarly to Ent5 (Hirst et al., 2003; Mills et al., 2003; Hirst et al., 2015). It is likely that many aspects of yeast Golgi protein recycling have counterparts in other eukaryotes.

## Materials and Methods

### Reagents and Tools

Table S1 lists the chemicals, primers, plasmids, and software used in this study, with details about source and availability.

### Yeast Growth and Transformation

The parental haploid *S. cerevisiae* strain was a derivative of JK9-3da (*leu2-3,112 ura3-52 rme1 trp1 his4*) (Kunz et al., 1993) carrying *pdr1Δ* and *pdr3Δ* mutations to facilitate HaloTag labeling (Barrero et al., 2016). Yeast cells were grown in baffled flasks with shaking at 23°C in the nonfluorescent minimal glucose dropout medium NSD (Bevis et al., 2002) or in the rich glucose medium YPD supplemented with adenine and uracil.

Yeast proteins were tagged by gene replacement using the pop-in/pop-out method to maintain endogenous expression levels (Rothstein, 1991; Rossanese et al., 1999). Plasmid construction was simulated and recorded using SnapGene. The plasmids generated in this study have been submitted to Addgene.

Deletion of *PDR1* and *PDR3* was accomplished by replacement with a G418 or nourseothricin resistance cassette from pFA6a-kanMX6 (Bähler et al., 1998) or pAG25 (Goldstein and McCusker, 1999), respectively. Deletion of *APL4* was accomplished by using overlap extension PCR to amplify a hygromycin resistance cassette from pAG32 (Goldstein and McCusker, 1999) flanked by 500 bp upstream and downstream of the gene. Deletion of *ENT5* was accomplished using the same overlap extension PCR strategy, except that the *LEU2* gene from *K. lactis* was amplified from pUG73 (Gueldener et al., 2002).

### CK-666 Treatment

Endocytosis was inhibited by incubating cells for the indicated time periods with 100 µM CK-666 diluted from a 50 mM stock solution in DMSO. Mock treatments employed DMSO alone.

### Fluorescence Microscopy

Live-cell 4D confocal microscopy was performed as previously described (Johnson and Glick, 2019). Yeast strains were grown in NSD and imaged at room temperature (∼23°C). Cells were attached to a concanavalin A-coated coverglass-bottom dish containing NSD, and were viewed on a Leica SP8 or SP5 confocal microscope equipped with a 1.4 NA/63X oil objective using a 60- to 80-nm pixel size, a 0.25- to 0.30-µm Z-step interval, and 20 to 30 optical sections. For time-lapse imaging, Z-stacks were captured at intervals of 2 s.

Static images and videos were deconvolved with Huygens Essential software using the classic maximum likelihood estimation algorithm (Day et al., 2017). With the aid of ImageJ (Schneider et al., 2012), images and videos were converted to hyperstacks and average projected, then range-adjusted to maximize contrast. Previously described custom ImageJ plugins were used to generate montages of time series, select individual structures and remove extraneous signal, convert edited montages to hyperstacks, and measure fluorescence intensities (Johnson and Glick, 2019). A new custom ImageJ plugin was used to average fluorescence traces, as described below.

### FM 4-64 Labeling and Quantification

For FM 4-64 labeling, a log-phase yeast culture was incubated for a 3- to 5-min pulse period with 0.8 µM FM 4-64FX diluted from a 1 mM stock in DMSO. Where indicated, external fluorescence was then quenched by addition of SCAS to a final concentration of 50 µM. Fluorescence was imaged on a Leica SP8 confocal microscope.

The quantification in Figure 1E was performed using ImageJ. For each sample, a confocal Z-stack was average projected, and then a mask representing the TGN cisternae was created using the Sec7-GFP signal. About 10% of the TGN cisternae could not be fully resolved from FM 4-64-containing non-TGN structures such as PVE compartments, vacuoles, and sites of polarized secretion, so those TGN cisternae were manually removed from the mask. The FM 4-64 signal within the adjusted mask was quantified for each cell, and this value was corrected by subtracting a background signal obtained from cell-free areas of the image.

### HaloTag Labeling

HaloTag labeling was performed as previously described (Casler et al., 2019). To visualize proteins fused to HaloTag, JFX_646_ or JFX_650_ ligand (Grimm et al., 2021), kindly provided by Luke Lavis (Janelia Research Campus, Ashburn, VA), was diluted 1:1000 from a 1 mM stock in DMSO in 0.5 mL of culture medium to give a final concentration of 1 µm. The medium was cleared of any precipitate by spinning at 17,000x*g* (13,000 rpm) in a microcentrifuge for 1 min. Then the cleared medium containing ligand was added to 0.5 mL of log-phase yeast culture, and the cells were incubated with shaking at 23°C for 30 min. Excess dye was removed by filtration through and washing on a 0.22-µm syringe filter (Millipore, catalog # SLGV004SL). The washed cells were resuspended in NSD and attached to a concanavalin A-coated coverglass-bottom dish. Videos were captured immediately by confocal microscopy.

### Statistical Analysis

Calculation and plotting of mean and SEM values was performed using Prism software. Each figure legend indicates the number of data points and experimental replicates.

### Quantification and Averaging of Golgi Maturation Events

Two- and three-color 4D confocal microscopy traces of individual cisternae containing tagged Golgi proteins were analyzed and quantified using custom ImageJ plugins as previously described (Johnson and Glick, 2019). For a single cisterna, the fluorescence trace in each channel was normalized to the average of the three highest values. For smoothing and averaging, a new custom ImageJ plugin was written to perform the following operations.

The first step was to smooth each noisy trace. For this purpose, a trace was numerically integrated, and then the integral was numerically differentiated using the smoothing method of Pavel Holoborodko (http://www.holoborodko.com/pavel/numerical-methods/numerical-derivative/smooth-low-noise-differentiators/) with n = 2, N = 11. The numerical differentiation formula is as follows. If f_0_ represents a given point in the integral, f_1_ represents the next point, f_-1_ represents the previous point, and so on, then the derivative at the point is estimated as

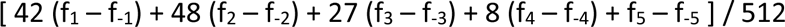

The peak of a smoothed trace was defined as the 50% point of the integral. To estimate the start and end points of a smoothed trace, a line was defined by the two time points when the trace first rose above 10% of its maximum value, and a second line was defined by the two time points when the trace permanently sank below 10% of its maximum value, and then these lines were extrapolated to the horizontal (time) axis.

The next step was to average the smoothed traces. For a given fluorescence channel, each smoothed trace was normalized to a maximum value of 1.0, and then the smoothed traces were averaged by centering at the peak values. The averaged trace was normalized to a maximum value of 1.0. To show multiple fluorescence channels on the same plot, the offset between any two fluorescence channels was calculated by measuring for each individual cisterna the offset between the peak values for the smoothed traces in the two channels, and then averaging those offset values. Plots of averaged traces show solid lines for mean values, plus shaded areas for 95% confidence intervals calculated by determining the SEM at each point and multiplying by 1.96. Relative arrival and departure times for two tagged Golgi proteins were calculated by measuring for each individual cisterna the offsets in start and end points for the smoothed traces in the two fluorescence channels, and then averaging those offset values.

The ImageJ plugins used for this study are available from a GitHub repository: https://github.com/bsglicker/4D-Image-Analysis.

## Acknowledgments

This work in the B.S.G. laboratory was supported by NIH grant R01 GM137004. J.C.C., A.H.K., and K.J.D. were supported by NIH training grant T32 GM007183. Thanks for assistance with fluorescence microscopy to Vytas Bindokas and Christine Labno at the Integrated Microscopy Core Facility, which is supported by the NIH-funded Cancer Center Support Grant P30 CA014599. We thank Luke Lavis for providing the JFX_646_ and JFX_650_ dyes.

---

**Figure S1.**
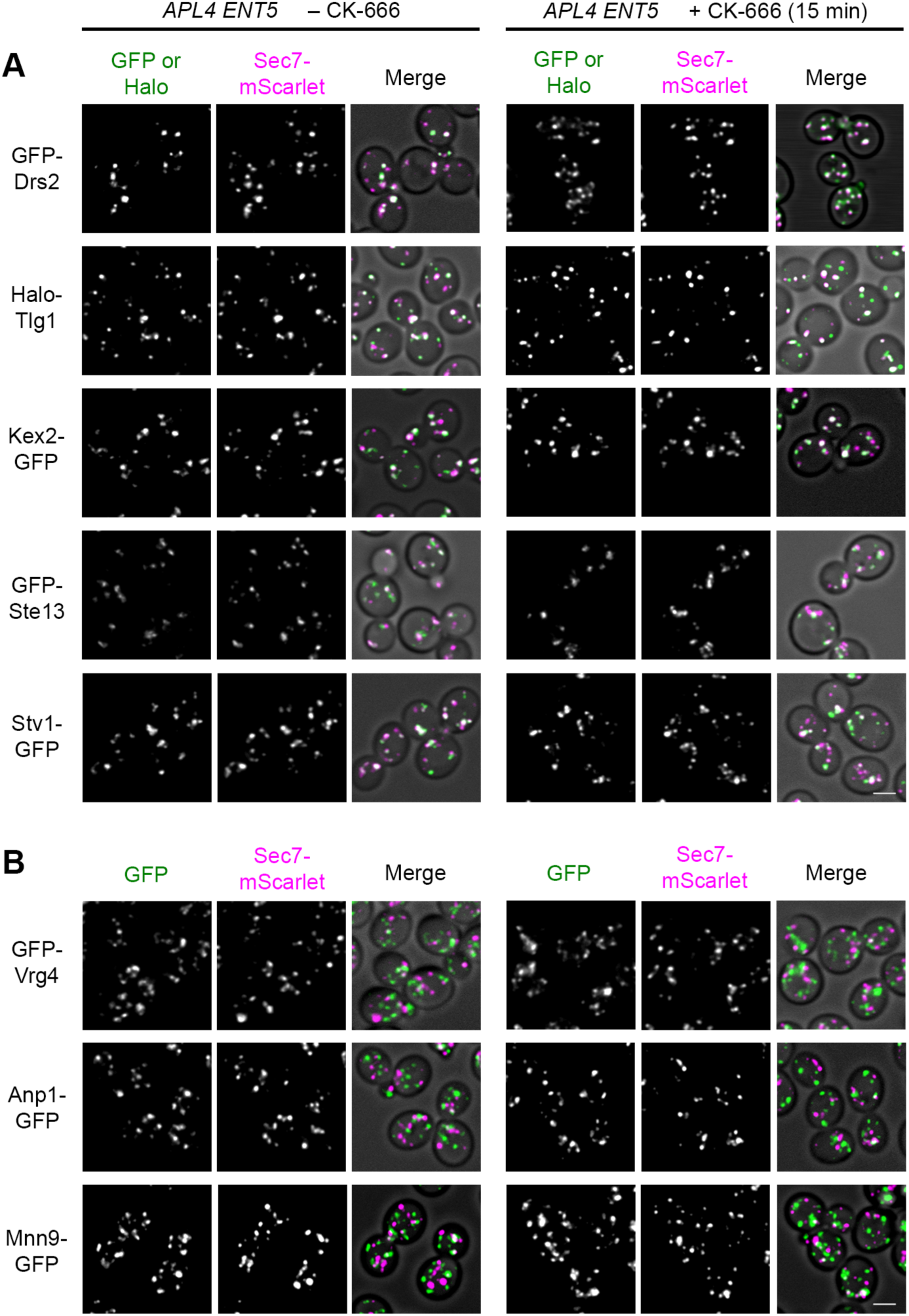
CK-666 Does Not Affect the Localizations of Transmembrane Golgi Proteins in Wild-Type Cells, Related to Figure 3. (A) Lack of a CK-666 effect on the localizations of transmembrane TGN proteins. The experiment was performed as in Figure 3D, except that *APL4 ENT5* cells were used and GFP-Ste13 and Stv1-GFP data are also shown. Scale bar, 2 µm. (B) Lack of a CK-666 effect on the localizations of transmembrane early Golgi proteins. The experiment was performed as in Figure 3D, except that *APL4 ENT5* cells were used and Anp1-GFP and Mnn9-GFP data are also shown. Scale bar, 2 µm.

**Figure S2.**
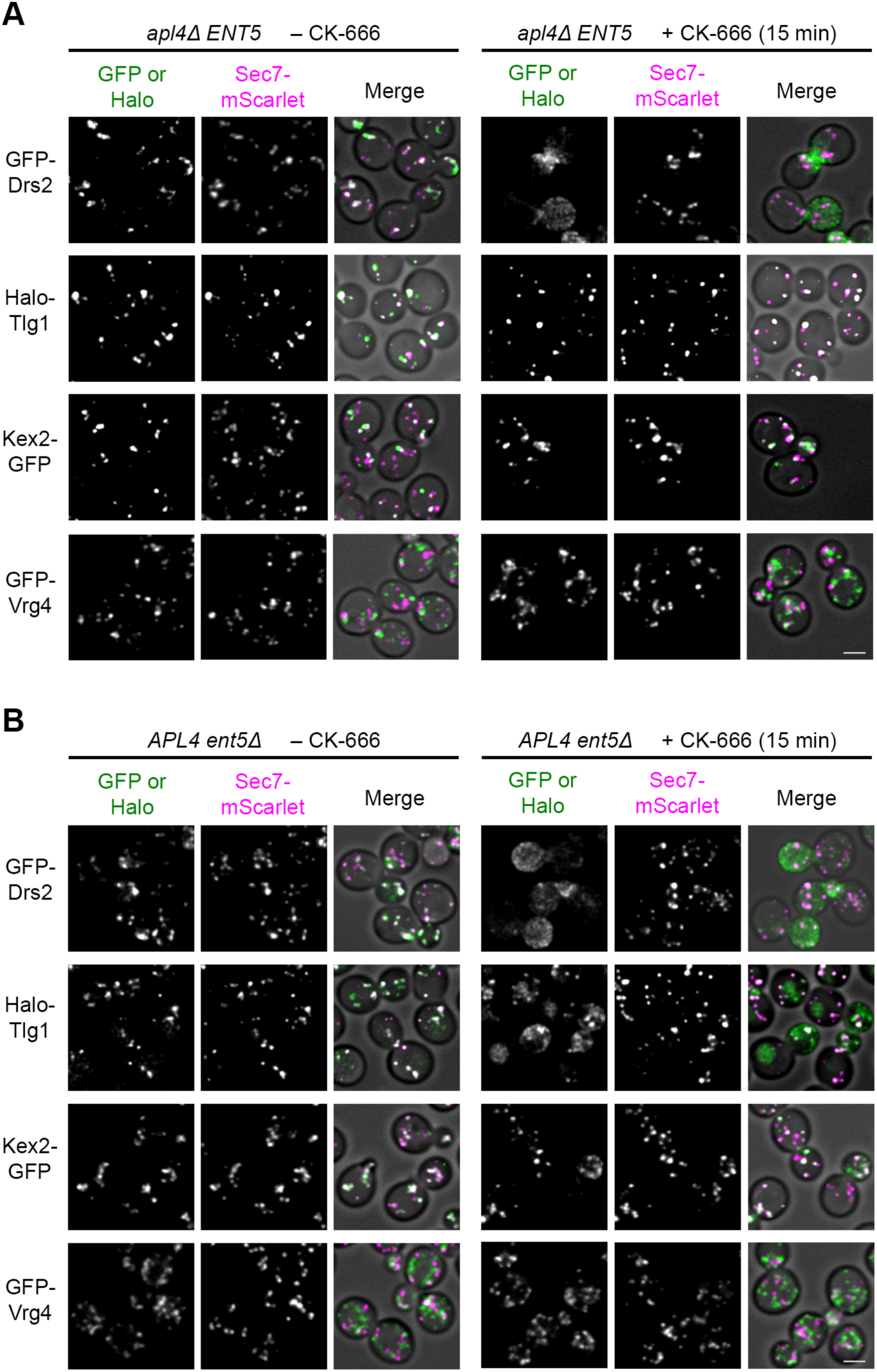
CK-666 Affects the Localizations of Some but Not All Transmembrane TGN Proteins in Single Mutant Cells Lacking Either AP-1 or Ent5, Related to Figure 3. (A) Effect of CK-666 on the localizations of transmembrane TGN proteins in the absence of AP-1. The experiment was performed as in Figure 3D, except that *apl4Δ ENT5* cells were used. CK-666 caused mislocalization of Drs2 but not of Tlg1, Kex2, or Vrg4. Scale bar, 2 µm. (B) Effect of CK-666 on the localizations of transmembrane TGN proteins in the absence of Ent5. The experiment was performed as in Figure 3D, except that *APL4 ent5Δ* cells were used. CK-666 caused mislocalization of Drs2 and Tlg1 but not of Kex2 or Vrg4. Scale bar, 2 µm.

**Figure S3.**
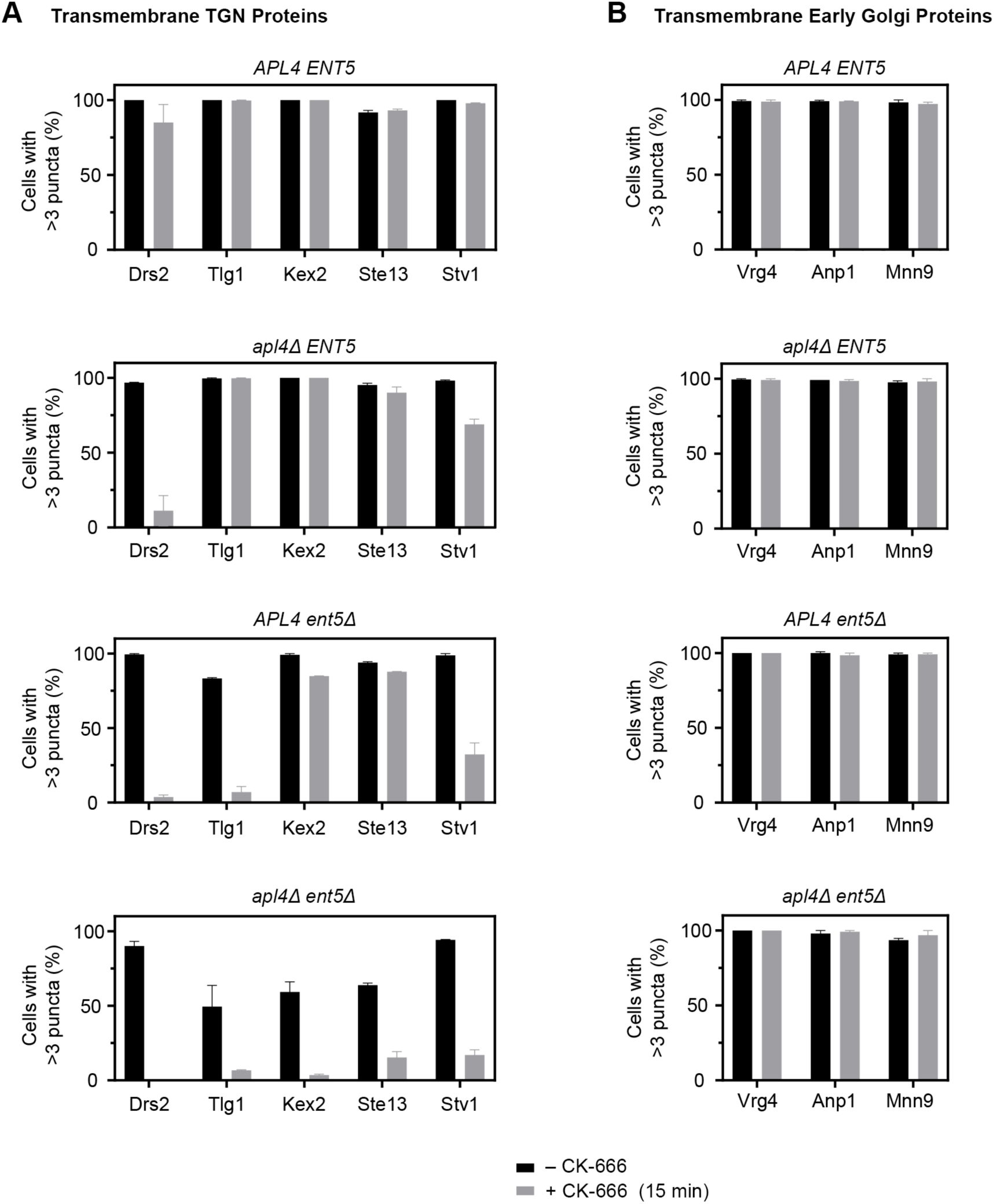
CK-666 Affects the Localization of Transmembrane TGN Proteins but Not of Transmembrane Early Golgi Proteins in Double Mutant Cells Lacking Both AP-1 and Ent5, Related to Figure 3. This figure extends Figure 3E, with the inclusion of additional marker proteins and of data from *apl4Δ ENT5* and *APL4 ent5Δ* cells. (A) Quantification of the CK-666 effects for five transmembrane TGN proteins in wild-type cells and in cells lacking either AP-1 or Ent5 or both. The experiment was performed as in Figure 3E. (B) Quantification of the CK-666 effects for three transmembrane early Golgi proteins in wild-type cells and in cells lacking either AP-1 or Ent5 or both. The experiment was performed as in Figure 3E.

**Figure S4.**
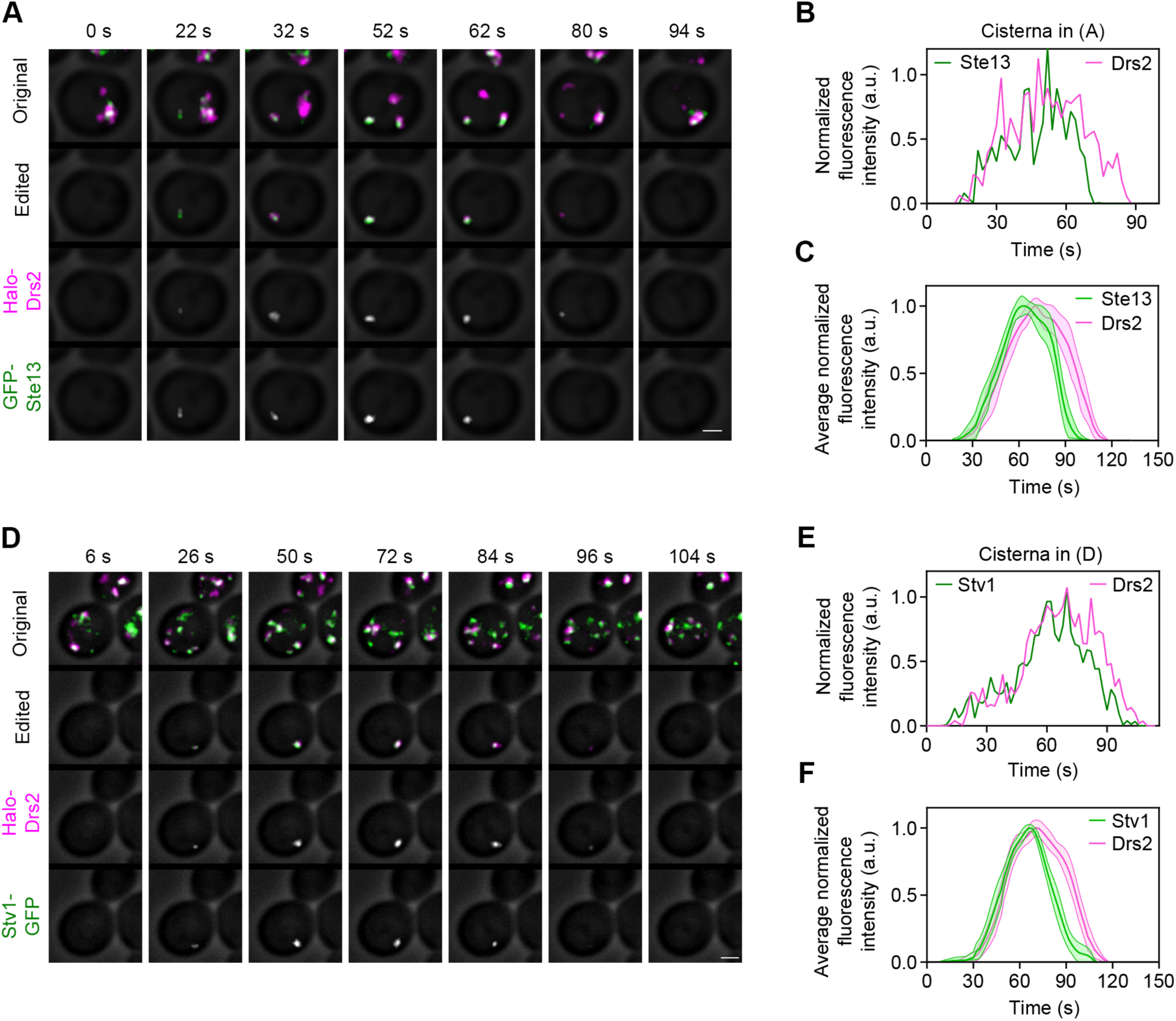
Additional TGN Proteins Resemble Drs2 in Their Kinetic Signatures, Related to Figure 4. (A) Maturation kinetics of Drs2 compared to Ste13. A strain expressing HaloTag-Drs2 and the dipeptidyl aminopeptidase GFP-Ste13 was grown to mid-log phase, labeled with JFX dye, and imaged by 4D confocal microscopy. Shown are average projected Z-stacks at the indicated time points from Part 3 of Video S4. The upper row shows the complete projections, the second row shows edited projections that include only the cisterna being tracked, and the subsequent rows show the individual fluorescence channels from the edited projections. Scale bar, 2 µm. (B) Quantification of tagged Golgi proteins during a typical maturation event. Depicted are the normalized fluorescence intensities in arbitrary units (a.u.) for the cisterna tracked in (A). (C) Smoothed and averaged traces showing the relative kinetic signatures of Drs2 and Ste13. Data were obtained for 10 representative cisternae. (D) Maturation kinetics of Drs2 compared to Stv1. The experiment was performed as in (A), except that the strain expressed the proton-pumping ATPase subunit Stv1-GFP, and the images are from Part 4 of Video S4. Scale bar, 2 µm. (E) Quantification of tagged Golgi proteins during a typical maturation event. Depicted are the normalized fluorescence intensities in arbitrary units (a.u.) for the cisterna tracked in (D). (F) Smoothed and averaged traces showing the relative kinetic signatures of Drs2 and Stv1. Data were obtained for 15 representative cisternae.

**Figure S5.**
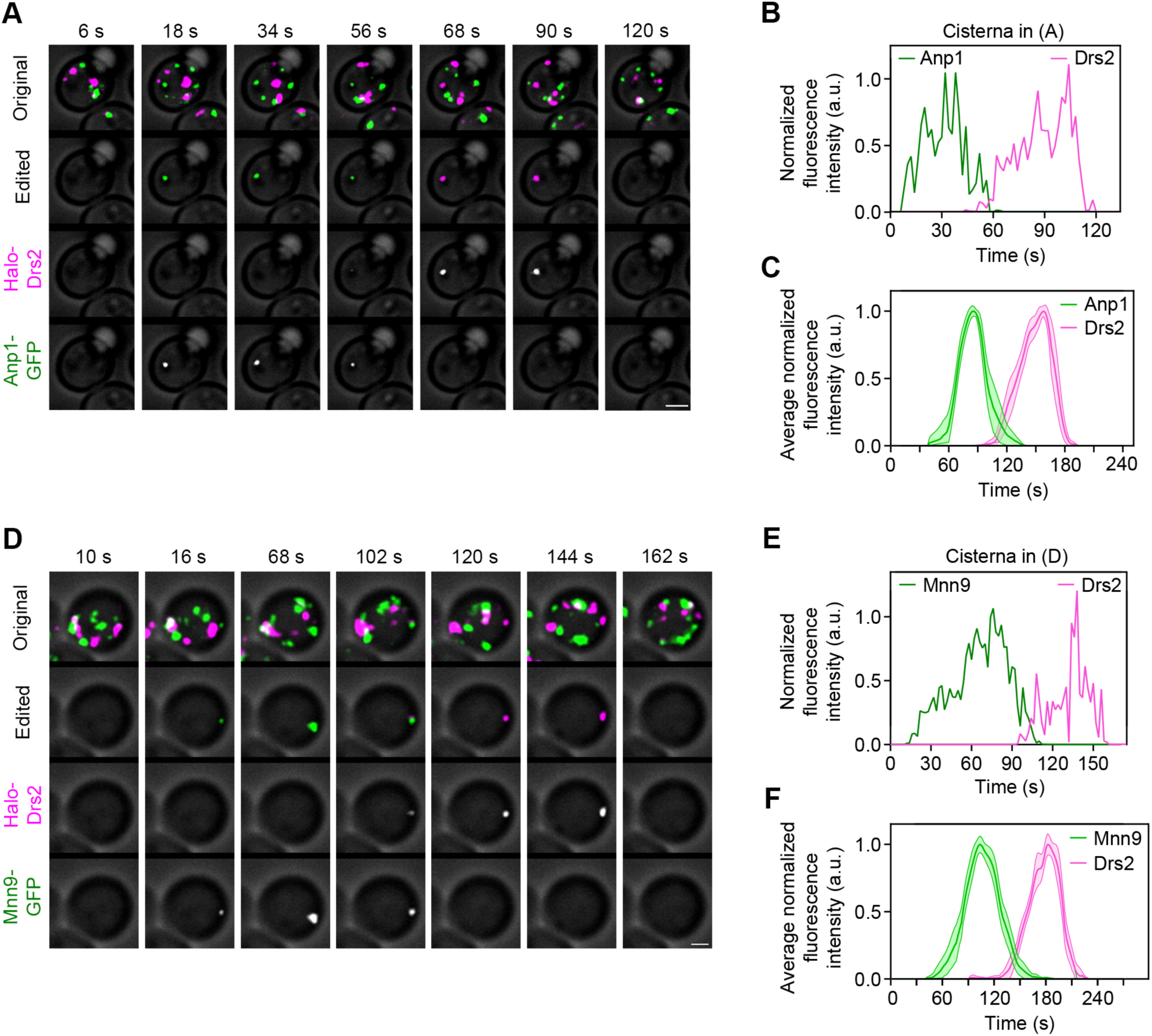
Additional Early Golgi Proteins Resemble Vrg4 in Their Kinetic Signatures, Related to Figure 4. (A) Maturation kinetics of Drs2 compared to Anp1. A strain expressing HaloTag-Drs2 and the mannosyltransferase Anp1-GFP was grown to mid-log phase, labeled with JFX dye, and imaged by 4D confocal microscopy. Shown are average projected Z-stacks at the indicated time points from Part 2 of Video S5. The upper row shows the complete projections, the second row shows edited projections that include only the cisterna being tracked, and the subsequent rows show the individual fluorescence channels from the edited projections. Scale bar, 2 µm. (B) Quantification of tagged Golgi proteins during a typical maturation event. Depicted are the normalized fluorescence intensities in arbitrary units (a.u.) for the cisterna tracked in (A). (C) Smoothed and averaged traces showing the relative kinetic signatures of Drs2 and Anp1. Data were obtained for 10 representative cisternae. (D) Maturation kinetics of Drs2 compared to Mnn9. The experiment was performed as in (A), except that the strain expressed the mannosyltransferase Mnn9-GFP, and the images are from Part 3 of Video S5. Scale bar, 2 µm. (E) Quantification of tagged Golgi proteins during a typical maturation event. Depicted are the normalized fluorescence intensities in arbitrary units (a.u.) for the cisterna tracked in (D). (F) Smoothed and averaged traces showing the relative kinetic signatures of Drs2 and Mnn9. Data were obtained for 13 representative cisternae.

**Figure S6.**
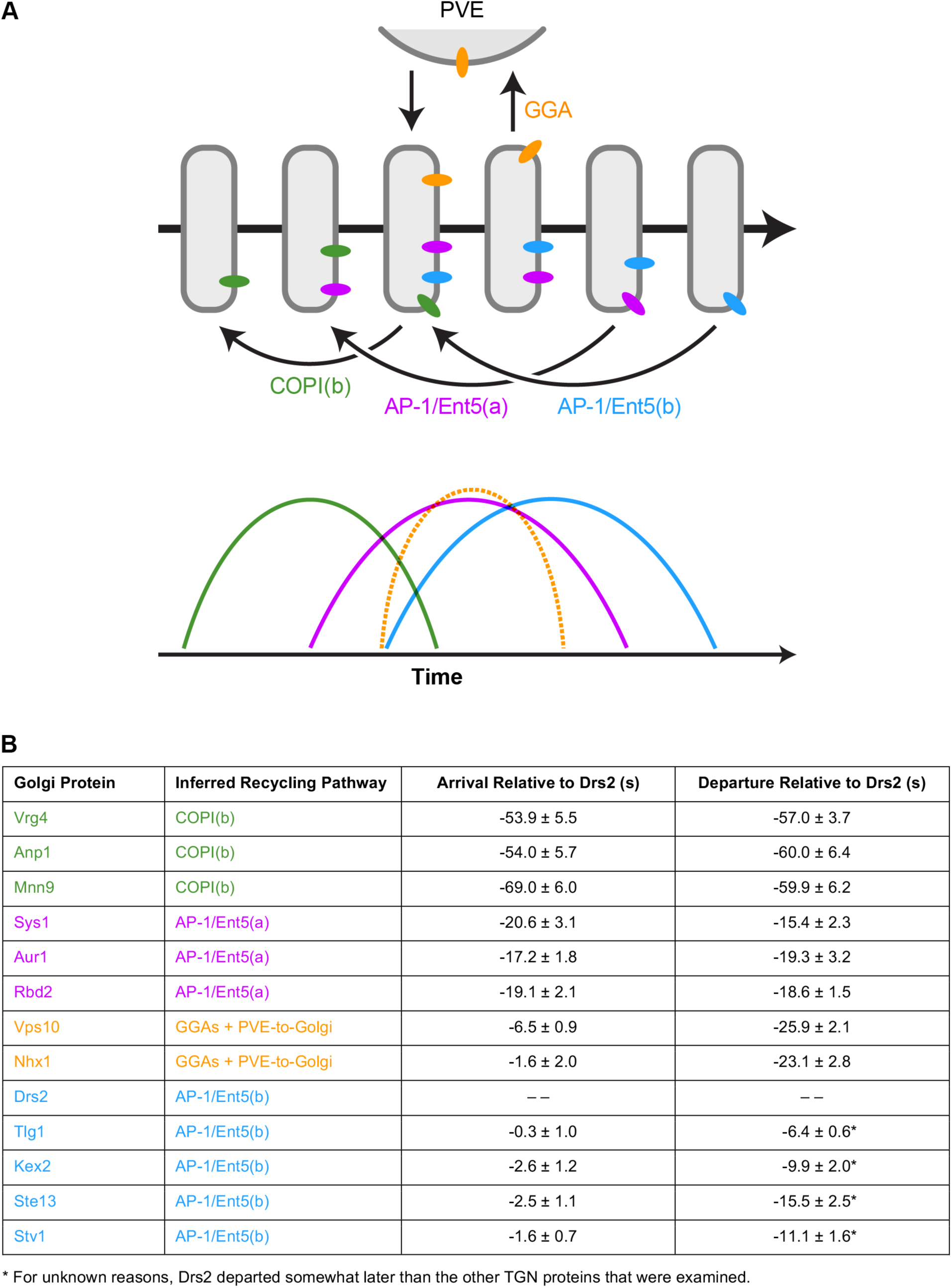
Recycling Pathways Determine the Kinetic Signatures and Localizations of Transmembrane Golgi Proteins, Related to Figures 1-7. (A) Model of Golgi recycling pathways and corresponding kinetic signatures. Straight horizontal arrows represent time during the life cycle of a cisterna, other arrows represent trafficking pathways, rounded rectangles represent Golgi cisternae of different ages, and colored ovals represent transmembrane proteins. Based on evidence presented here and elsewhere, we suggest that there are three sequential intra-Golgi recycling pathways. The first pathway is designated COPI(b) to distinguish it from COPI(a)-mediated Golgi-to-ER recycling, which is not depicted. This COPI(b) pathway generates early kinetic signatures (green). The second pathway is designated AP-1/Ent5(a), and it recycles proteins such as Sys1 to generate intermediate kinetic signatures (magenta). The third pathway is designated AP-1/Ent5(b), and it recycles TGN proteins such Drs2 to generate late kinetic signatures (blue). In addition, a pathway of GGA-dependent transport to prevacuolar endosome (PVE) compartments followed by recycling to nascent TGN cisternae generates separate kinetic signatures (orange). For a given transmembrane Golgi protein, the kinetic signature governs localization to cisternae in a particular age range. Some Golgi proteins might follow more than one pathway, in which case their kinetic signatures would be combinations of the ones depicted here. (B) Quantification of kinetic signatures for Golgi transmembrane proteins that are believed to follow four distinct recycling pathways. The colors match those in (A). Shown are average and SEM values for arrival and departure times relative to Drs2, based on the same data that were used to generate plots in other figures. Negative numbers indicate arrival or departure earlier than Drs2.

**Video S1. AP-1 and Ent5 Are Present Late in TGN Maturation, Related to Figure 1**

A strain expressing the TGN marker Sec7-mScarlet, the AP-1 subunit Apl2-GFP, and the clathrin adaptor Ent5-HaloTag was imaged as described in Figure 1A. Shown are average projected Z-stacks. Time is indicated in min:sec format. The upper panel shows the complete projections, the second panel shows edited projections that include only a single representative cisterna, and the subsequent panels show the individual fluorescence channels from the edited projections. Scale bar, 2 µm.

**Video S2. Golgi Maturation Is Perturbed by CK-666 in a Strain Lacking AP-1 and Ent5, Related to Figure 2**

Scale bars, 2 µm.

(Part 1) Normal Golgi maturation in an untreated *apl4Δ ent5Δ* cell. A strain expressing the early Golgi marker GFP-Vrg4 and the TGN marker Sec7-mScarlet was imaged as described in Figure 2D. Shown are average projected Z-stacks. Time is indicated in min:sec format. The upper panel shows the complete projections, the second panel shows edited projections that include only a single representative cisterna, and the lower two panels show the individual fluorescence channels from the edited projections.

(Parts 2 and 3) Abnormal Golgi maturation in *apl4Δ ent5Δ* cells. The experiment was performed as in Part 1, except that the cells were treated with CK-666 to block endocytosis as described in Figure 2F. In each case, the upper panel shows the complete projections and the lower panel shows edited projections that include only a single representative cisterna, which persistently contained either the early Golgi marker (Part 2) or the TGN marker (Part 3).

**Video S3. Drs2 Departure Begins at About the Same Time as AP-1 Arrival, Related to Figure 3**

A strain expressing the TGN marker Sec7-mScarlet, the AP-1 subunit Apl2-GFP, and HaloTag-Drs2 was imaged as described in Figure 3A. Shown are average projected Z-stacks. Time is indicated in min:sec format. The upper panel shows the complete projections, the second panel shows edited projections that include only two representative cisternae, and the subsequent panels show the individual fluorescence channels from the edited projections. Scale bar, 2 µm.

**Video S4. Four Transmembrane TGN Proteins Arrive and Depart at About the Same Time as Drs2, Related to Figure 4**

Each part of the video has the following format. Average projected Z-stacks are shown. Time is indicated in min:sec format. The upper panel shows the complete projections, the second panel shows edited projections that include only a single representative cisterna, and the lower two panels show the individual fluorescence channels from the edited projections. Scale bars, 2 µm.

(Part 1) A strain expressing GFP-Drs2 and HaloTag-Tlg1 was imaged as described in Figure 4A.

(Part 2) A strain expressing HaloTag-Drs2 and Kex2-GFP was imaged as described in Figure 4D.

(Part 3) A strain expressing HaloTag-Drs2 and GFP-Ste13 was imaged as described in Figure S4A.

(Part 4) A strain expressing HaloTag-Drs2 and Stv1-GFP was imaged as described in Figure S4D.

**Video S5. Three Early Golgi Proteins Arrive and Depart Much Earlier than Drs2, Related to Figure 4**

Each part of the video has the following format. Average projected Z-stacks are shown. Time is indicated in min:sec format. The upper panel shows the complete projections, the second panel shows edited projections that include only a single representative cisterna, and the lower two panels show the individual fluorescence channels from the edited projections. Scale bars, 2 µm.

(Part 1) A strain expressing HaloTag-Drs2 and GFP-Vrg4 was imaged as described in Figure 4G.

(Part 2) A strain expressing HaloTag-Drs2 and Anp1-GFP was imaged as described in Figure S5A.

(Part 3) A strain expressing HaloTag-Drs2 and Mnn9-GFP was imaged as described in Figure S5D.

**Video S6. Proteins that Travel from the TGN to PVE Compartments Depart Earlier than Drs2, Related to Figure 5**

Each part of the video has the following format. Average projected Z-stacks are shown. Time is indicated in min:sec format. The upper panel shows the complete projections, the second panel shows edited projections that include only a single representative cisterna, and the lower two panels show the individual fluorescence channels from the edited projections. Scale bars, 2 µm.

(Part 1) A strain expressing HaloTag-Drs2 and Vps10-GFP was imaged as described in Figure 5A.

(Part 2) A strain expressing GFP-Drs2 and Nhx1-HaloTag was imaged as described in Figure 5D.

**Video S7. The Transmembrane Golgi Protein Sys1 Has an Intermediate Kinetic Signature, Related to Figure 6**

Each part of the video has the following format. Average projected Z-stacks are shown. Time is indicated in min:sec format. The upper panel shows the complete projections, the second panel shows edited projections that include only a single representative cisterna, and the subsequent panels show the individual fluorescence channels from the edited projections. Scale bars, 2 µm.

(Part 1) A strain expressing Sec7-mScarlet, HaloTag-Drs2, and Sys1-GFP was imaged as described in Figure 6A.

(Part 2) An *apl4Δ ent5Δ* strain expressing Sec7-mScarlet and Sys1-GFP was imaged as described in Figure 6F.

**Video S8. Two Transmembrane Golgi Proteins Arrive and Depart at About the Same Time as Sys1, Related to Figure 7**

Each part of the video has the following format. Average projected Z-stacks are shown. Time is indicated in min:sec format. The upper panel shows the complete projections, the second panel shows edited projections that include only a single representative cisterna, and the subsequent panels show the individual fluorescence channels from the edited projections. Scale bars, 2 µm.

(Part 1) A strain expressing HaloTag-Drs2 and Aur1-GFP was imaged as described in Figure 7C.

(Part 2) A strain expressing HaloTag-Drs2 and Rbd2-GFP was imaged as described in Figure 7F.

**Table S1.**
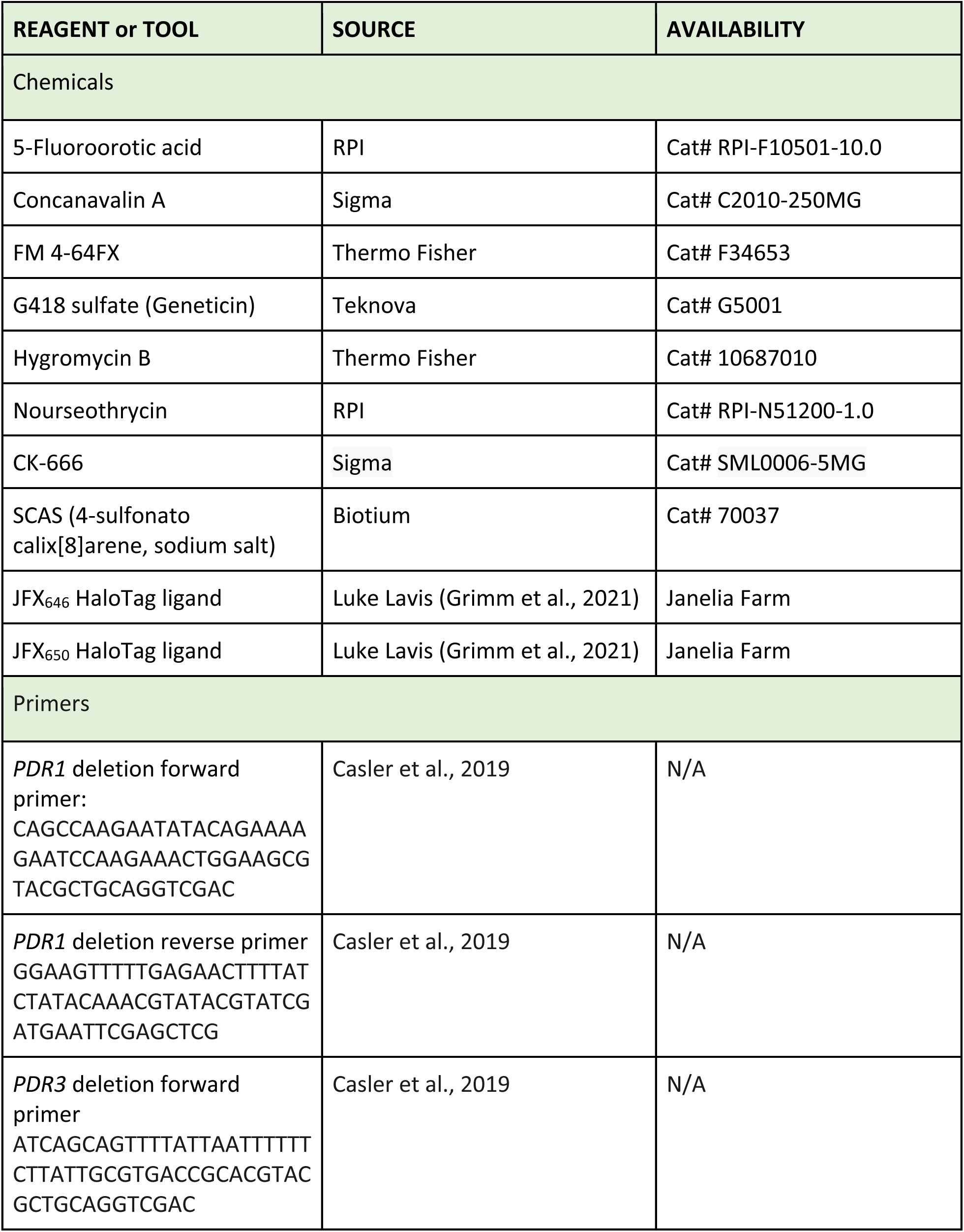
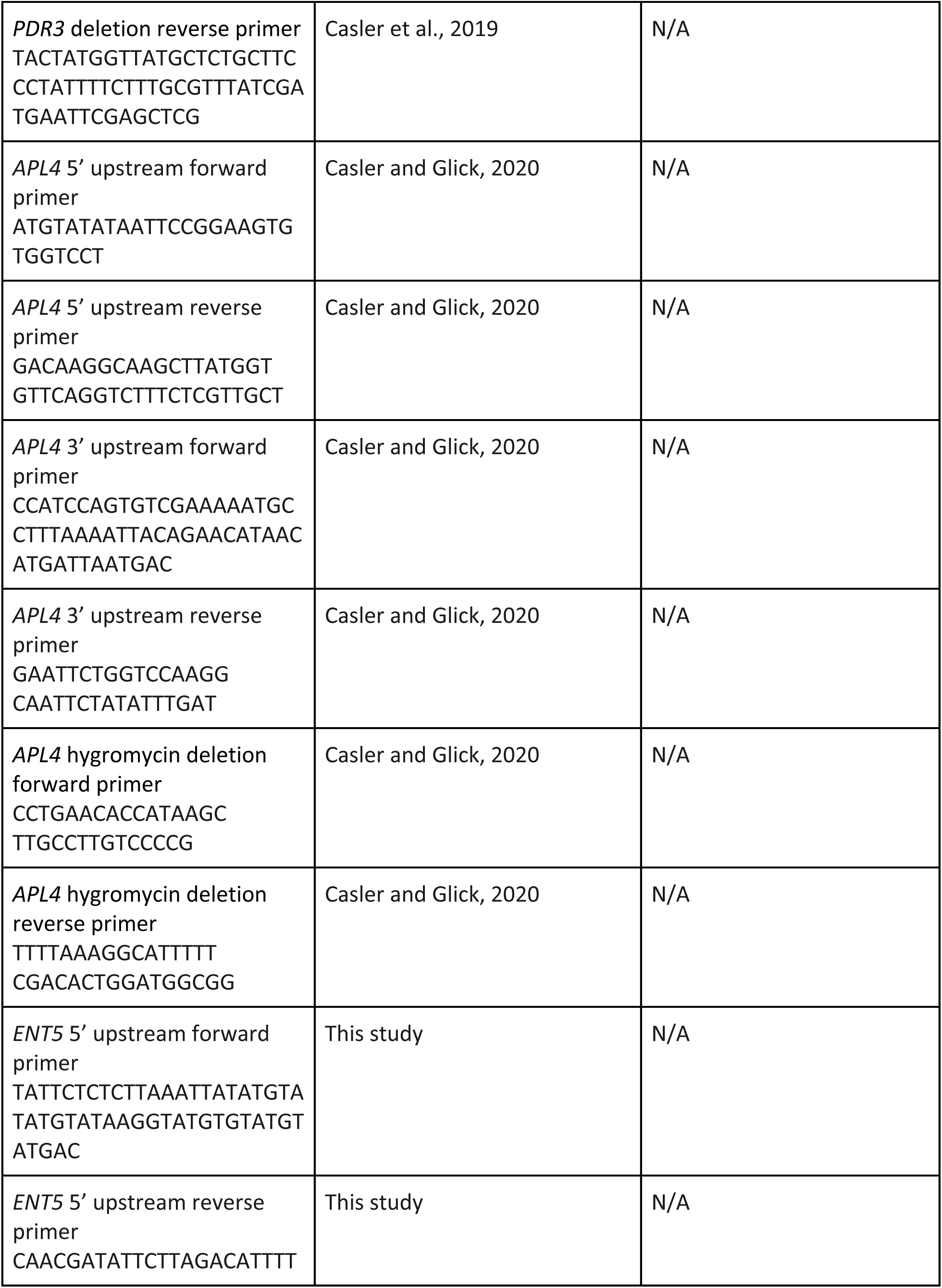
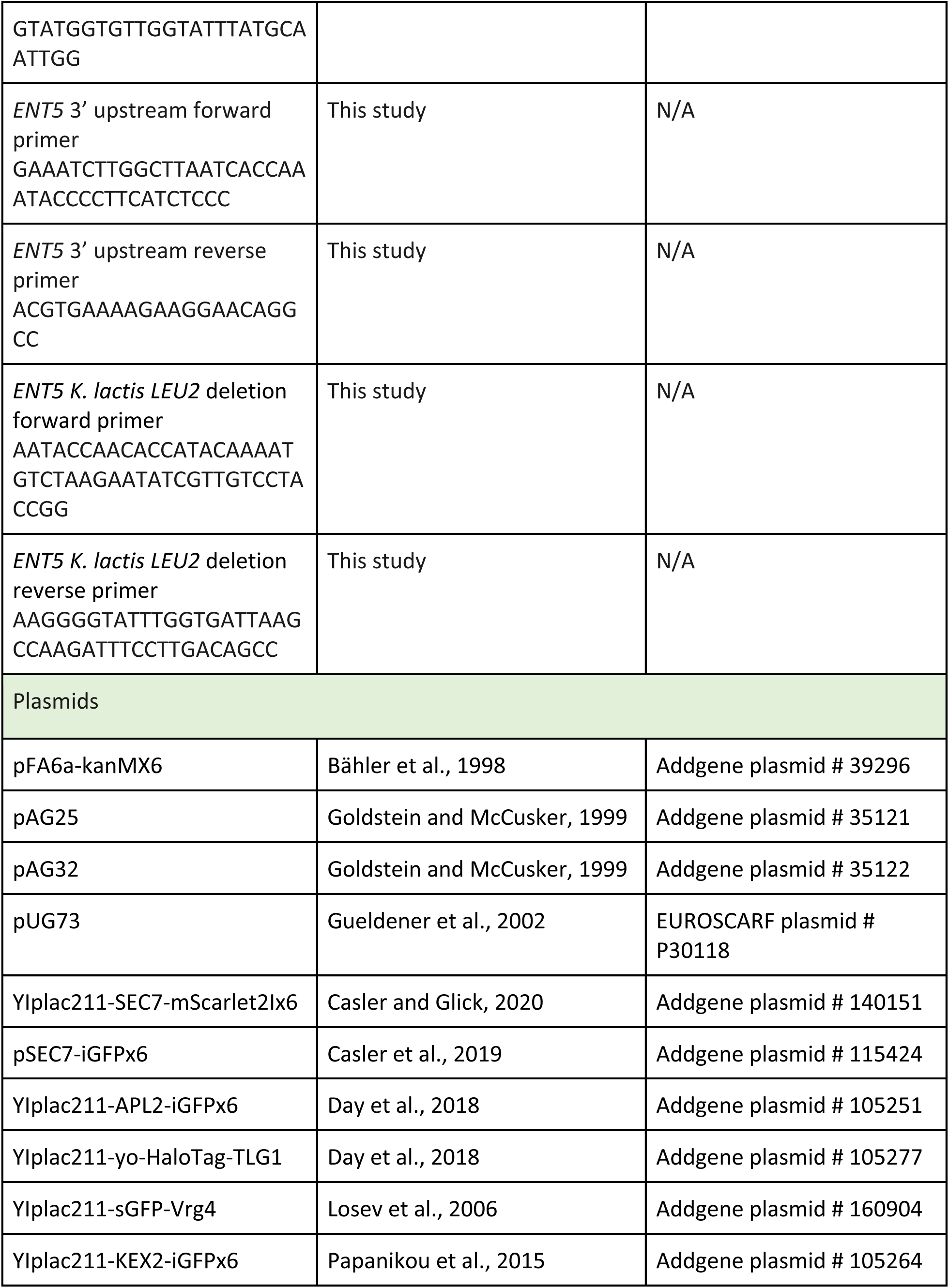
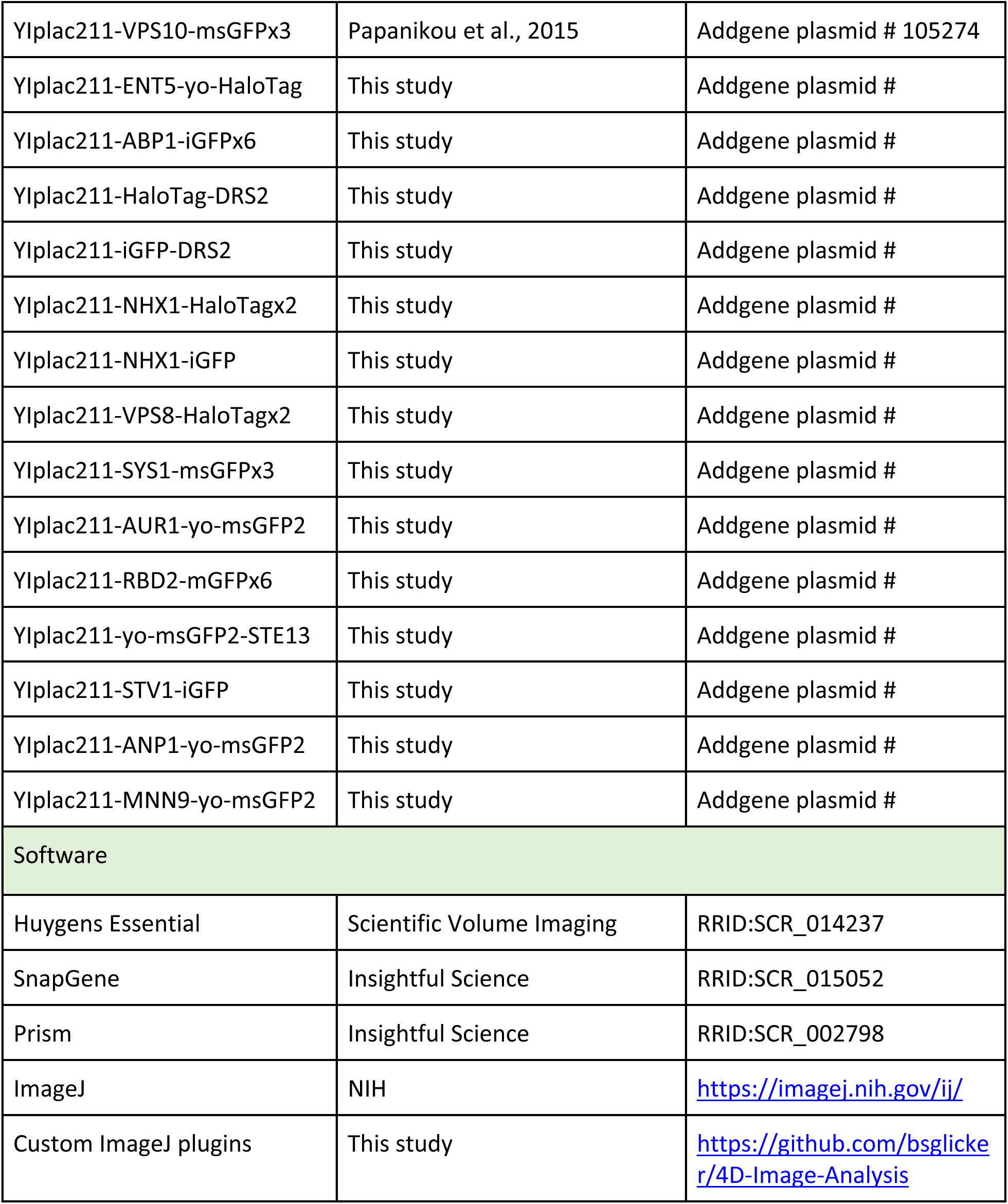
Reagents and tools used in this study.

